# SpIC3D imaging: Spinal In-situ Contrast 3D imaging

**DOI:** 10.1101/2025.03.05.641747

**Authors:** Lucy Liang, Alessandro Fasse, Arianna Damiani, Maria K. Jantz, Uzoma Agbor, Matteo del Brocco, Rachel Monney, Cecelia Rowland, Taylor Newton, Chaitanya Gopinath, David J. Schaeffer, Christi L. Kolarcik, Lee E. Fisher, Esra Neufeld, Marco Capogrosso, Robert A. Gaunt, T. Kevin Hitchens, Elvira Pirondini

## Abstract

High-definition visualization techniques are critical for understanding the neuroanatomy of the spinal cord, an essential structure for sensorimotor and autonomic functions, in both healthy and pathological conditions. Magnetic resonance imaging (MRI) is a common method for visualizing neural structures in 3D. However, techniques for spinal cord MRI have historically achieved limited visualization of rootlets and nerves, especially at lower spinal levels, due to their highly complex and compact organization. Here we developed a spinal in-situ contrast 3D imaging (SpIC3D) method that allows visualization of spinal compartments in fixed animal and human specimens with ultra-high resolution at various spinal levels. Using SpIC3D, we achieved quantification of neuronal cell density in dorsal root ganglia, multi-segment identification of individual rootlets and roots, and volumetric reconstruction of multiple spinal structures for computational modeling. SpIC3D provides a basis for accelerated spinal pathology characterization and personalized spinal cord stimulation treatments.

## INTRODUCTION

The spinal cord is an essential part of the central nervous system for the control of both sensorimotor and autonomic functions. It connects the brain to the rest of the body and is encapsulated within the vertebral column, surrounded by a layer of epidural fat, meninges, and cerebral spinal fluid (CSF) ^1,2^. It gives rise to several nerve root pairs, exiting superior or inferior to the corresponding vertebra with decreasing root angles from rostral to caudal spine areas ^3^. Analysis of white matter fibers and alterations in whole anatomical structures is critical to understand spinal function and dysfunction. Specifically, whole-spine high-resolution imaging would allow quantitative analysis of spinal structures, cell distributions, root and rootlet fiber orientations, and their alterations in pathological conditions ^4^. Additionally, a high-resolution template of the spine structures and fiber anatomy could enhance neurosurgical approaches, implantation procedures, and functional interpretations ^5–9^.

High-resolution postmortem magnetic resonance imaging (hrMRI) of fixed tissues has recently emerged and shows high potential towards these aims. hrMRI can overcome drawbacks from more common approaches such as histological staining and neuronal tracing, which are traditional but labor-intensive processes, used to analyze cellular distributions and to trace axonal projections. Indeed, histological steps such as sectioning and subsequent 3D reconstruction can lead to significant tissue loss and deformation when applied to small and complex structures like rootlet and root geometries along the spinal cord. Additionally, while recent advancements in whole-tissue imaging technology based on light-sheet fluorescence microscopy coupled with tissue clearing methods, such as CLARITY ^10^, CUBIC ^11,12^, PACT and PARS ^13^, ScaleS ^14^, and iDISCO+ ^15^, have partially enabled intact 3D volume visualization, these techniques are limited to small tissue volumes and suffer from clearing-induced morphological changes, hampering accurate geometric measurements ^16,17^. MRI, instead, can resolve 3D anatomy and connectivity in much larger samples, at unprecedented resolution, and with minimal distortion.

While a promising technique, there are still many challenges associated with common postmortem hrMRI approaches. Many groups attempted spinal cord postmortem hrMRI in rats ^18–22^, cats ^23–25^, monkeys ^26,27^, and humans ^4,28,29^. However, so far, because of hardware limitations, scan time costs, and bone-soft tissue interface artifacts, the spinal cord was removed from the vertebral column for imaging, consequently disturbing epidural fat, the CSF-containing dural sac, as well as root and rootlet fibers, impeding the visualization of the spine structures and the spinal nerves’ natural anatomical organization, a limitation also affecting histological and clearing processes.

If this organization could be preserved, high-resolution anatomical MRI and diffusion-weighted MRI (dMRI) could be leveraged to map the root fiber pathways. However, while diffusion imaging and tractography have been extensively studied in the brain ^30–32^, optimization of dMRI acquisition protocols for the spinal cord is still understudied. Generally, axon fibers in the brain radiate and cross in all directions around a spherical shape. Spinal roots, instead, are highly organized and unidirectional, mainly running along the rostrocaudal axis, challenging our ability to differentiate across multiple nerves, particularly in the more caudal areas. This substantially different anatomical organization undermines the translation of brain dMRI approaches to the spinal cord and roots. The majority of previous efforts in adapting dMRI for the spinal cord were in-vivo and focused on the cervical spinal cord to quantify fractional anisotropy (FA) and mean diffusivity (MD) or detect spinal cord pathologies, such as tumors and demyelination ^33–38^. Few selected studies used tractography ^39–46^, with very few looking at scan optimization, and the nerve roots and spinal cord were often examined separately. Out of these, none examined the organization of spinal roots within the spinal column, the primary targets of recent neuromodulation approaches—such as spinal cord stimulation for sensory and motor restoration — as well as of many common surgical procedures ^47–49^. Therefore, volumetric images and visual differentiation of spinal roots at the lumbosacral level using tractography in an intact spinal column still represents a major challenge.

In this study, we developed a novel spinal in-situ contrast 3D imaging method (SpIC3D) that achieves imaging of the full spinal column at ultra-high resolution. It includes multiple sequences and contrasts to visualize different spinal structures and quantify neuronal cell density, as well as an optimized diffusion weighted imaging protocol for reconstructing and differentiating spinal roots and rootlets. Additionally, with SpIC3D we established the first-ever highly realistic multi-tissue computational lumbosacral spinal cord model of electrical stimulation exclusively based on imaging data free from structural distortions. Importantly, we demonstrated that SpIC3D can generalize to different spinal levels and scale up to the human spinal cord. This method has the benefit of combining the high resolution of postmortem imaging and the intact structure of in-vivo imaging, providing a great tool for constructing a comprehensive and detailed spinal atlas.

## RESULTS

### SpIC3D acquisition and preparation pipeline optimization

We developed a novel spinal in-situ contrast 3D imaging (SpIC3D) pipeline that allows 3D reconstruction of anatomical structures at ultra-high-resolution (50 µm) in cat and human spines (**Figure 1A-B**). Unlike previous accounts of spinal cord ultra-high-resolution imaging, the bone structure was kept intact, preserving the surrounding 3D features of the spinal cord (**Figure 1C**). Several factors contributed to our ability to obtain these ultra-high-resolution data.

**Figure 1.**
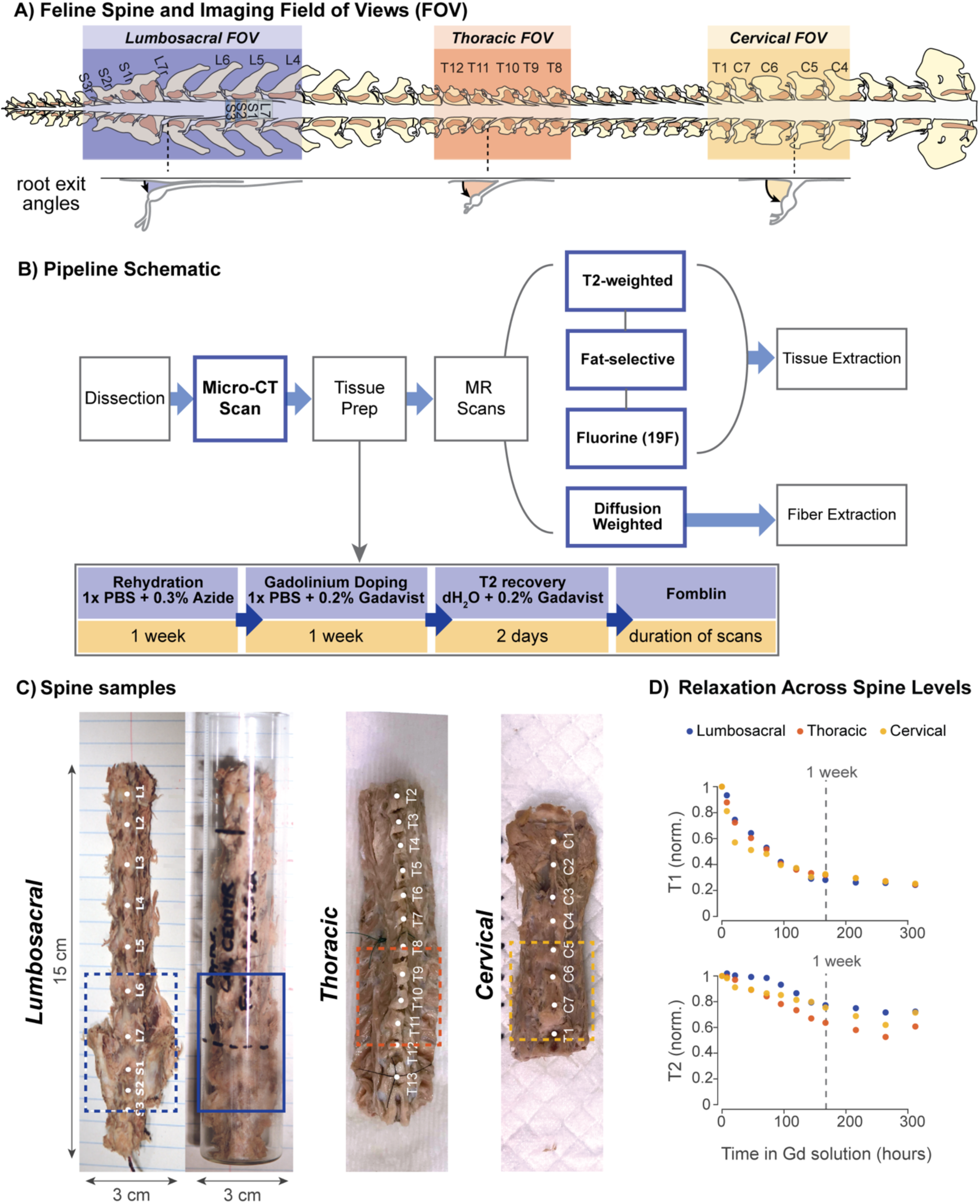
Sample preparation and image acquisition pipeline. **(A)** Cervical, thoracic, and lumbosacral spinal levels commonly of interest for spinal cord stimulation (and thus imaging targets in this study), represented in the cat spinal cord. Enlarged visualizations of representative roots from all spine regions show the substantial decrease in root-spinal cord angle from cervical to lumbosacral roots. **(B)** Flow diagram showing the full tissue preparation, multimodal imaging, and segmentation pipeline. The five acquired imaging sequences are highlighted with blue frames. **(C)** *Left:* dissected cat lumbosacral spine sample (C01-LS) with dimensions, and sample placed tightly in a 30 mm OD glass tube that fits in the RF coil. The 4 cm imaging field of view (FOV) is demarked by a blue rectangle. *Middle*: dissected cat thoracic spine sample (C05-Th), with 4 cm FOV (red rectangle). *Right*: dissected cat cervical spine sample (C05-Ce), with 4 cm FOV (yellow rectangle). **(D)** Normalized T1 (*top*) and T2 (*bottom*) relaxation time changes in 0.2% Gd over time for one lumbosacral, one thoracic, and one cervical sample. The reduction rate is consistent across spinal regions. T1 relaxation is reduced by 70-80% and stabilizes after around 1 week (dotted line), while T2 continues to shorten, but at a much slower rate.

A T1 relaxation-optimized tissue preparation protocol was fundamental to implementing the high-resolution multi-shell diffusion protocol. Following tissue isolation and high-resolution CT scanning for bone segmentation, the specimens were soaked in a 0.2% solution of Gadavist (1.0 M Gadobutrol). This gadolinium-based T1 relaxation agent can significantly shorten the tissue water T1 and has been shown to facilitate 3D high-resolution MRI and multi-shell diffusion MRI (dMRI) ^50^. We monitored the mean tissue T1 and T2 relaxation times periodically during the preparation period and determined that one week is sufficient to reduce the T1 by 70-80% while the T2 is reduced by only about 20-30% (**Figure 1D, Figure S1**). This reduction allowed us to use rapid MR imaging protocols with a short recycle delay (TR: repetition time) between excitation pulses. Importantly, the reduction rates were consistent across spinal regions, samples, and species supporting the generalizability of our protocol (**Figure 1D, Figure S1**). Following the tissue preparation, the sample was submersed in Fomblin, an inert perfluorocarbon oil that provides magnetic susceptibility matching with a 1H-signal-free background and protects the sample from dehydration (**Figure 2A**).

**Figure 2.**
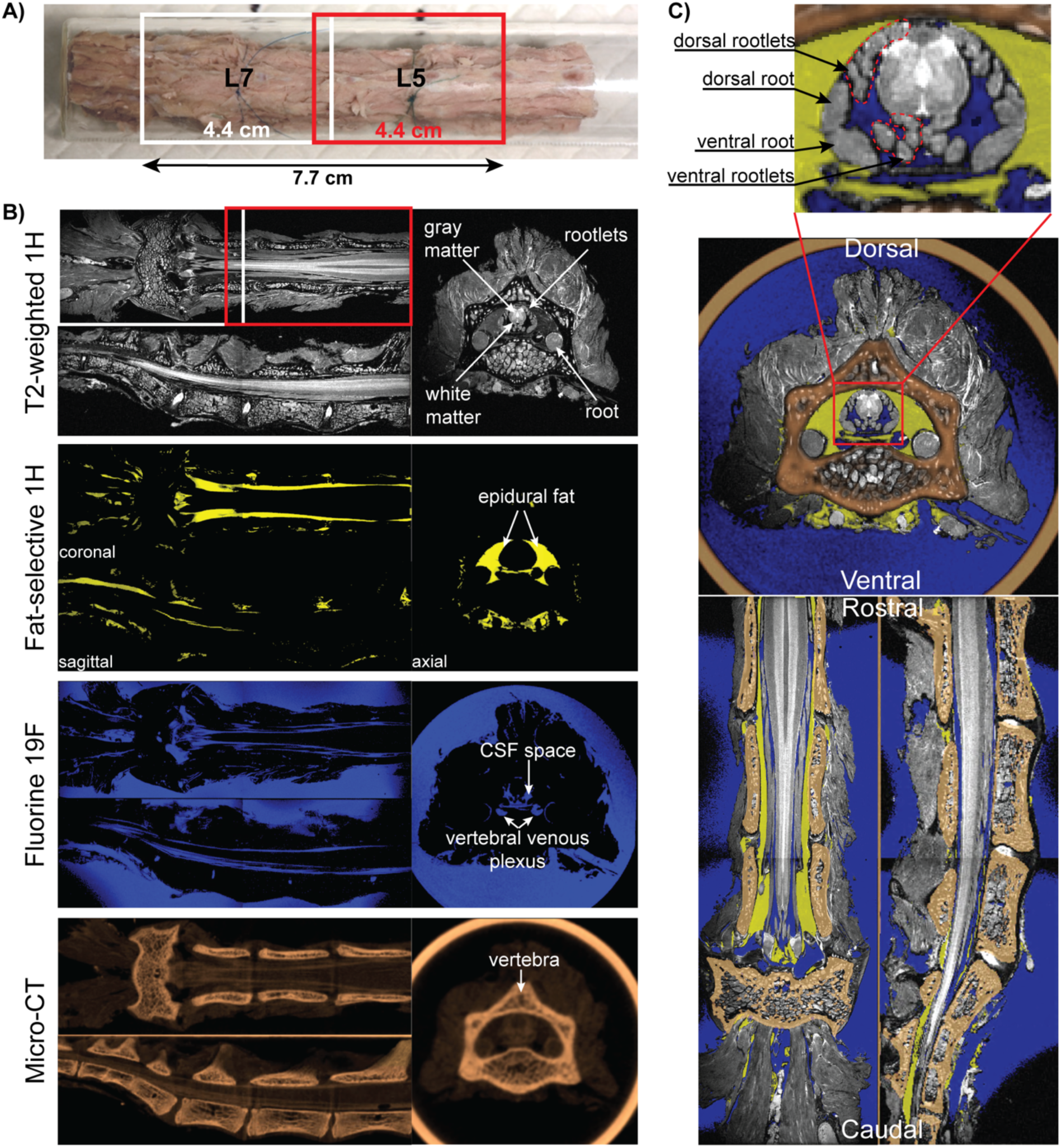
Multi-modal imaging of the cat lumbosacral spinal cord. **(A)** Cat lumbosacral spine sample (C03-LS) in 3 cm glass tube. Two overlapping 4.4 cm imaging FOVs are shown as white and red rectangles, centered on the L7 and L5 vertebrae, respectively, resulting in a final 7.7 cm long image of the spine after merging. **(B)** Merged and registered images (50 µm in-plane, 100 µm inter-slice resolution) shown for three anatomical planes and the four acquisition modalities: T2-weighted (grayscale), fat-selective (yellow), 19F MRI for Fomblin (blue) MR images, and a micro-CT (copper). The combination of these acquisitions provides clear visualization of different spinal structures and compartments. Importantly, also the vertebral venous plexus is visible in the images (see more details in **Figure S3**). **(C)** Merged image of all imaging modalities in B). Inset: clear visualization of ventral/dorsal roots and rootlets.

Another important factor was the cutting-edge gradient and radiofrequency (RF) technology implemented in high-field scanners. Specifically, the cat spinal cords were scanned at 11.7 T using a vertical-bore microimaging system with a Mirco2.5 gradient set capable of 1,500 mTm^−1^ and a Mini0.75 gradient set capable of 450 mTm^−1^, paired with a 30 mm or 40 mm inner diameter quadrature RF coil, respectively. Vertical positioning is helpful in sample preparation, but our approach generalizes to standard horizontal-bore rodent imaging systems and lower fields (i.e., 9.4 T/30 cm with a 12 cm high-performance gradient set and 72 mm RF coil). Importantly, spinal levels are typically much more rostral than the corresponding root and vertebral levels, particularly in the lumbosacral spine (**Figure 1A**). To extend the imaging beyond the RF coil field of view (FOV), we used a tiling technique. Specifically, a MINIWB57 probe with Animal Handling System that allows for graduated vertical translation of the sample in the probe was used to acquire 2 separate 40 mm FOV datasets with a small sample overlap, allowing for simple end-to-end registration (**Figure 2A**) with no additional computational manipulation.

With these optimizations, SpIC3D achieves ultra-high-resolution (50 µm in-plane, 50-100 µm slice thickness) and facilitates the distinction of spinal structures (outlined in **Figure 1B**) in acceptable time (111 +/- 14 hrs). In comparison, we estimated that an expert technician would require about 20 hours (∼3 days) to histologically pre-process (i.e., freeze, section, mount) a sample of comparable size (800 sections). It should be noted that the reported time for SpIC3D includes all sequences necessary to obtain anatomical details of multiple tissues. Histological processing, by contrast, still requires additional steps, including multiple laborious staining protocols (e.g., Nissl, H&E) as well as imaging, extending total process time. Most importantly, for SpIC3D, human involvement is only required during sample and scanner setup, and not all acquisitions need to be performed in one setting, which means that typically under-exploited scanner time, such as nights, weekends, and holidays, can be efficiently utilized.

### SpIC3D sequences capture accurate anatomy of the cat lumbosacral spine

SpIC3D includes a micro-CT acquisition (100 μm isotropic) to capture the bone structure and a T2-weighted 1H sequence to image white and gray matter (**Figure 2B**). Importantly, T2-weighted resolution is a tradeoff between the smallest structure of interest and acquisition time. Here we selected 50 µm resolution allowing us to visualize individual rootlets, which are typically 100-200 µm in cats, in 9.5 hrs (**Figure 2C**). Yet, it is possible for SpIC3D to achieve a higher resolution (25 µm) if desired, at the cost of a longer acquisition time (38 hrs, **Figure S2**), but such a resolution was not necessary for the cat spine. Additionally, we developed two novel sequences to visualize the epidural fat and the CSF, as well as large vessels of the vertebral venous plexus (**Figure 2B, Figure S3**). Epidural fat is an important tissue that serves both as a shock absorber and a lubricant for the spinal cord ^51^. While it is commonly selectively suppressed in T2-weighted 1H sequences to improve contrast to neural tissue, here we took advantage of its properties to develop a T2-weighted sequence that selectively enhances epidural fat. For the CSF, this is typically not present in a post-mortem tissue, especially after dissection. However, as demonstrated in our images (**Figure 2B**), the CSF cavity is preserved and filled with Fomblin. Using fluorine-sensitive 19F MRI, we were able to visualize the Fomblin and thus estimate the original CSF distribution. This method also reveals the large vertebral venous plexus vessels, which closely resembles those commonly visualized with contrast CT venography ^52^ (see example in **Figure S3A**), but with better resolution and vessel reconstruction (**Figure S3B-C**). Finally, high-resolution diffusion-tensor imaging was implemented to distinguish roots and rootlets at each spinal level. Once more, the visualization of these spinal compartments was possible because the vertebrae were preserved intact.

We optimized the spinal diffusion-weighted MRI (dMRI) protocol to determine the minimum angularity needed for accurate diffusion tensor reconstruction (**Figure 3**). While similar efforts have been extensively pursued for the brain, very few studies attempted angularity optimization for the spinal cord, which has a fundamentally different white matter organization, with the majority of fibers running parallel, eventually forming roots, and exiting the spinal canal with segment-specific angles (**Figure 1B**) ^34,36,38^. We began optimization by obtaining “ground truth” dMRI data that had not only the high spatial-resolution necessary for dissecting the different rootlets and roots, but also a high angular resolution to capture the organization of the dorsal root entry zones. For this, we acquired a dMRI dataset of sample C01-LS with 0.2 mm isotropic resolution and a multi-shell diffusion scheme with b-values 1000 s/mm² and 3500 s/mm² using 12 and 49 diffusion sampling directions, respectively. Directions were generated with the Emmanuel Caruyer online q-space sampling protocol application ^53^ (**Figure S4A**). We then simulated 55 subsets of diffusion data with a range of 6-61 directions uniformly distributed around a sphere, and for each subset we computed fiber tractography through common regions of interest (ROI) seeding at the L6, L7, S1, and S2 roots (**Figure 3B**). For all roots, quantitative measurements (fractional anisotropy [FA], mean diffusivity [MD], tract length, tract diameter, tract volume) showed a rapid change with the increase in number of diffusion sampling directions. The curves flatten at around 19 diffusion gradients (**Figure 3C, S4B**) suggesting this schema as sufficient for accurate 3D reconstruction of roots and rootlets. The 19-diffusion schema was sufficient to precisely reconstruct spinal nerve fibers as they travel from spinal roots to rootlets and into the spinal cord, and to automatically label individual rootlets within the spinal canal (**Figure 3B**). The diameter of L6, L7, S1, and S2 roots reconstructed using 19 directions were validated by in-vitro measurements (<200 µm error when comparing root diameter measured from images and dissection, **Figure 3D**). We therefore established a dMRI protocol with 19 diffusion sampling directions that were either over a single shell with b-value 2000 s/mm² (samples C03-LS and C04-Th) or split over 2 shells (b-values 1000 s/mm² and 2000 s/mm² or 3500 s/mm², samples C04-LS, C05 and C06) and we replicated it in three additional animals (**Figure S4A, Supplementary Table 1**). Importantly, no differences in fiber tract reconstruction were visually observable between single and 2 shell protocols.

**Figure 3.**
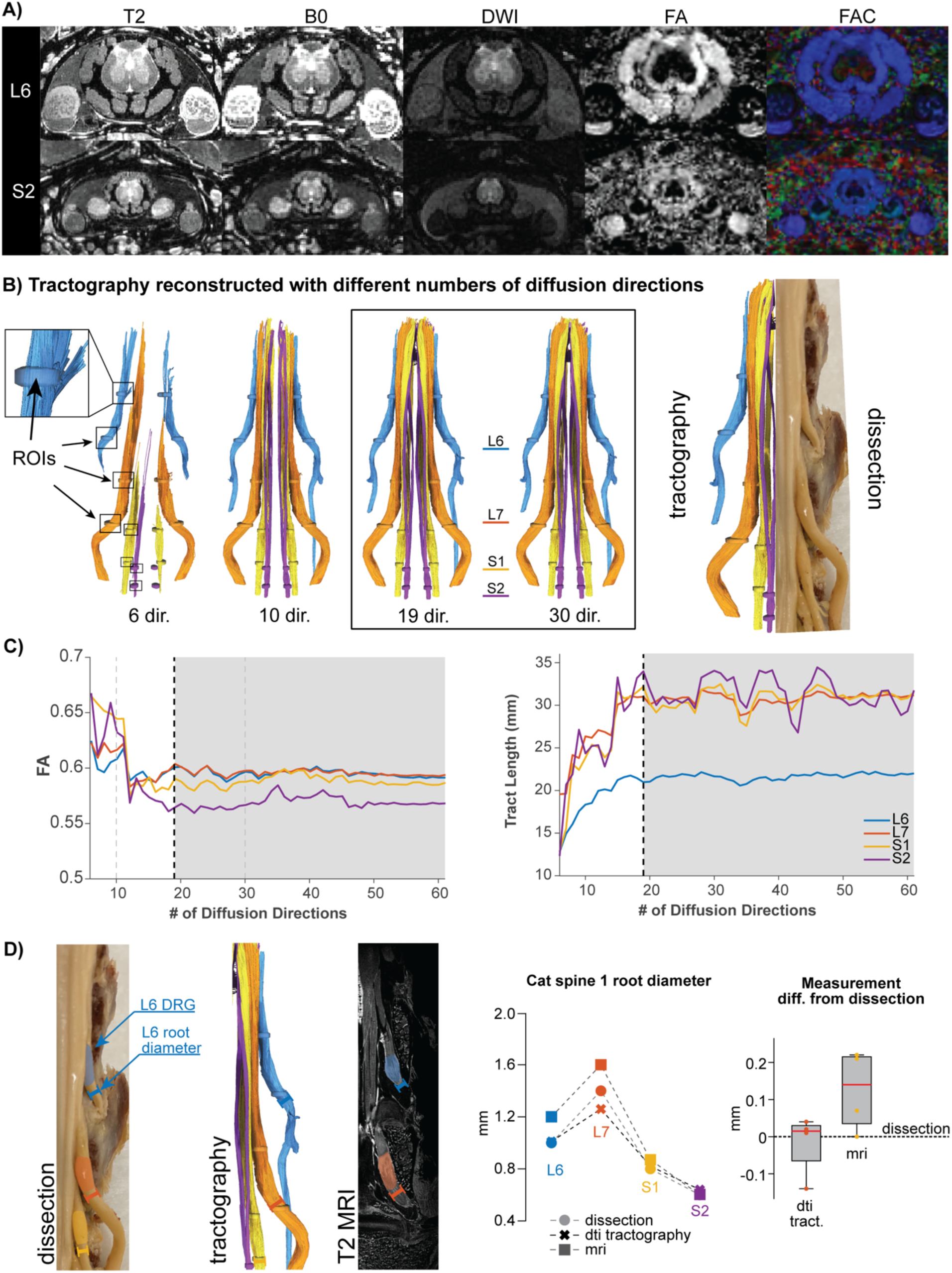
Diffusion tensor imaging protocol optimization and tractography validation. **(A)** Multi-contrast axial magnetic resonance microscopy images of the C01-LS spinal cord at L6 and S1 levels. Spinal cord levels are named based on the exiting spinal roots using standard abbreviations: L for lumbar and S for sacral. Contrasts include: *T2*, T2-weighted gradient echo (50 µm); *B0*, b=0 image from diffusion acquisition (200 µm); *DWI*, isotropic diffusion weighted image (200 µm); *FA*, fractional anisotropy (200 µm); *FAC*, directionally colored fractional anisotropy (200 µm). **(B)** *Left*: tractography results reconstructed using varying numbers of diffusion directions (6, 10, 19, 30 directions, respectively) from a diffusion weighted dataset with a total of 61 diffusion directions (b=1000/3500). The tracts are colored based on the root level they belong to. *Right*: comparison of tractography (19 direction) and dissection of the same cat spine sample. **(C)** Tractography quality as measured by FA and tract length stabilizes at 19 directions and above (gray). **(D)** *Left*: visual representation of root diameter (past the DRG) measurements based on tissue dissection, DTI tractography, and T2 MRI. *Right*: plot comparing root diameters measured with the different methods for each of the 4 root levels (L6, L7, S1, S2), and box plots of measurement errors (n=4 root levels) compared to ground truth measures from tissue dissection. For all boxplots, the whiskers extend to the maximum and minimum. Central line, top, and bottom of the box represent median, 75th percentile, and 25th percentile, respectively.

Overall, SpIC3D includes a collection of 1) a micro-CT to visualize the vertebrae, 2) an anatomical T2w image to visualize the ensemble of the cord, 3) a fat-selective image to capture epidural fat coverage, 4) a 19F image to visualize the CSF, and 5) a 19 direction DTI sequence to visualize detailed root fiber trajectories (**Supplementary Video 1**, parameters summarized in **Supplementary Table 1**). This innovative ensemble of sequences offers a unique basis for examining the finely-detailed 3D structures within the spinal canal and local fiber orientations within the white matter.

### SpIC3D allows quantification of spinal root entry zones and dorsal ganglia cell distributions

Each dorsal root is attached to the dorsolateral sulcus of the spinal cord by a series of rootlets arranged in a line – the dorsal root entry zone (DREZ). Similarly, ventral root fibers exit the cord forming an elliptical area named the anterior (ventral) root exit zone (AREZ). Dorsal rootlet fibers extend from the DREZ into the spinal gray matter and eventually “terminate” as the axon bundles, branch, disperse, and synapse onto spinal neurons. We will refer to this fiber penetration into the gray matter as gray matter entry zone. SpIC3D was not only able to reliably capture length and diameter of roots but also to visualize DREZ, AREZ, and gray matter entry zone. So far, quantification of grey matter entry zone was only possible via neuronal tracers and post-mortem histological analysis, which are cumbersome processes. Additionally, tracers of different colors are necessary to investigate the organization of different roots, limiting the number of fibers that can be studied.

Here we show that the ultra-high resolution of SpIC3D overcomes these limitations, allowing the visualization of different axons within the gray matter at different spinal levels (**Figure 4A**). Additionally, SpIC3D allowed us to reliably quantify the extent of DREZ and AREZ of different roots (**Figure 4B**). We found that left and right DREZ are not perfectly symmetric and may have some overlap between root levels. AREZs tend to be wider than the corresponding DREZ, and gray matter entry zones are rostral to the DREZ. We validated the DTI-measured DREZ length using measurements obtained from the dissected tissues (**Figure 4C**). Finally, SpIC3D also allowed us to resolve the sacral plexus, i.e., an intricate ensemble of nerves, which is formed by the sacral spinal roots S1-S3 and receives contributions from the L7 lumbar spinal root (**Figure 4D**). To further elucidate the innervation of pelvic, pudendal, and sciatic nerves within the plexus, for which proportion of contribution from each spinal root is still unclear, we applied SpIC3D to a wider and longer sample (C02) in which the three nerves were preserved intact during tissue preparation. For this, we first implemented SpIC3D in a larger bore scanner with standard quadrature coil (9.4T, **Figure S5**), demonstrating the valuable applicability of SpIC3D in more common horizontal acquisition set-ups. SpIC3D was able to precisely image and differentiate the three nerves and their complex organization over several roots (**Figure S5**). One can readily see that the sciatic nerve consists of fiber contributions from L6, L7, S1 spinal roots, and to a small extent from the S2 root. The pelvic nerve, on the other hand, only receives contributions from S1, S2, and S3, while the pudendal nerve is formed by spinal roots L7, S1, S2, and S3. These results are consistent with dissection studies in literature ^54–57^, further highlighting the potential of SpIC3D.

**Figure 4.**
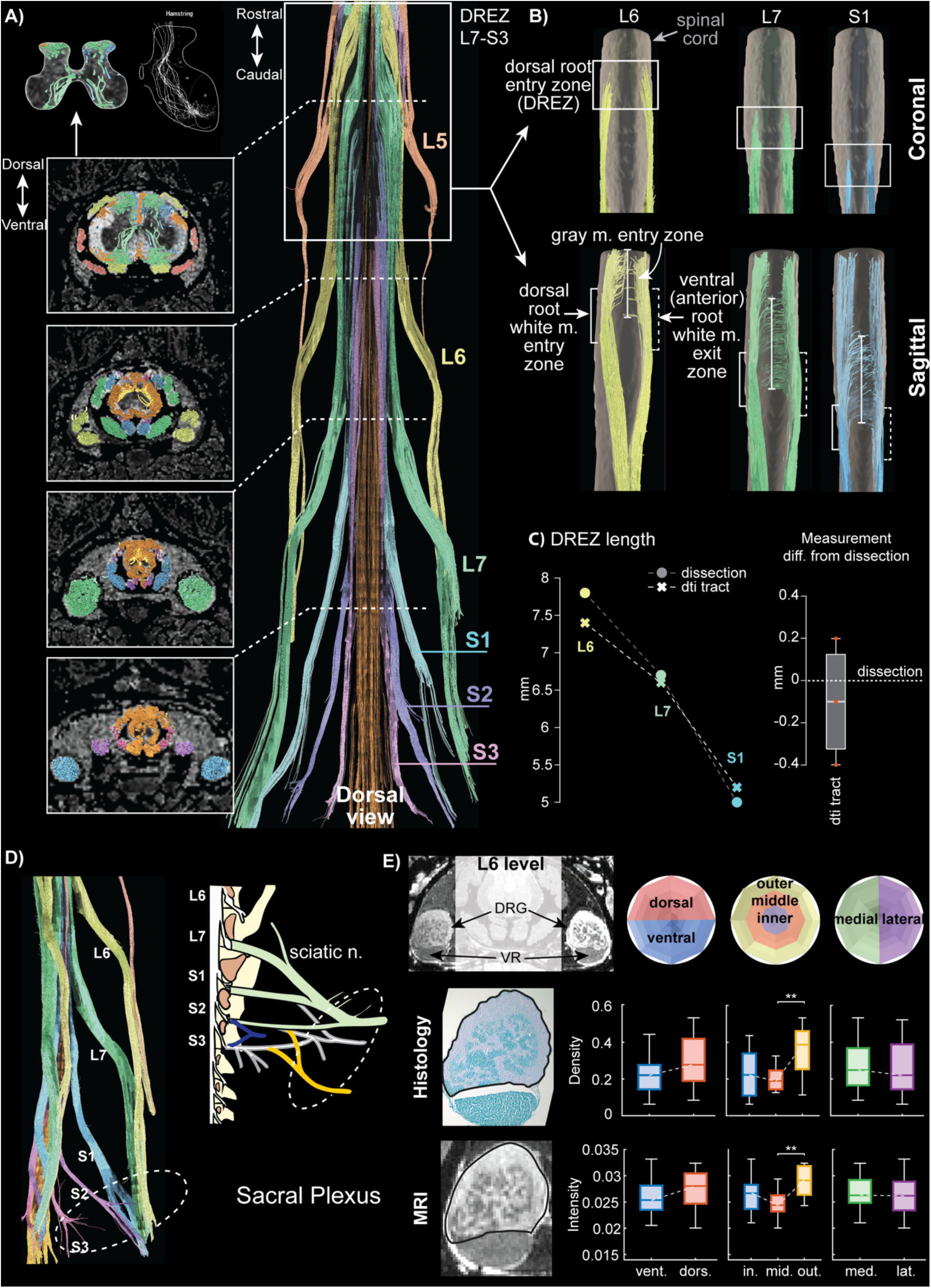
Detailed morphological features captured from DTI and T2-weighted MRI. **(A)** Sample C03-LS. *Left*: *Top* - intra-gray matter trajectories of the L7 root fibers resemble stained afferent fiber trajectories reported by Ishizuka et al., 1979 ^59^. *Bottom* - axial slices from each of the L5, L6, L7, S1 root levels show fiber distinction both in the rootlets and white matter. Caudal root fibers are more medial in the spinal gray matter than the rostral root fibers. *Right*: Tract reconstruction from seeds in regions-of-interest defined at each spinal root allows root-level-specific rootlets and spinal white matter fiber differentiation. **(B)** Dorsal root entry zones (DREZ), ventral (anterior) root exit zones (AREZ), as well as gray matter entry zones visualized with tractography for L6, L7, S1 root fibers. **(C)** *Left*: Plot comparison of DREZ lengths measured through DTI tractography and dissection at L6, L7, S1 levels, and box plot of differences between measures (n=3 spinal levels) from DTI and T2-weighted MRI and measures from dissected tissues (i.e., ground truth). **(D)** *Left*: Proximal portion of the sacral plexus visualized with tractography. *Right*: schema of the sacral plexus adapted from Nur et al., 2021 ^54^. **(E)** Quantification of cell distribution within the L6 DRG using T2-weighted MRI, validated against histological data from the same DRG sample, using a variant of the quantification method and algorithm from Ostrowski et al., 2017 ^60^. Boxplots from left to right show dorsal/ventral distribution (n=24 areas per group), inner/middle/outer distribution (n=16 areas per group), and medial/lateral distribution (n=24 areas per group). The outer ring cell density is significantly higher than the middle ring (Mann-Whitney U test for two groups or Kruskal-Wallis test with Bonferroni correction for three groups were used for significance comparisons, **: p<0.01, no significance p>0.05 if not indicated). For all boxplots, the whiskers extend to the maximum and minimum. Central line, top, and bottom of the box represent median, 75th percentile, and 25th percentile, respectively.

Importantly, dorsal and ventral roots fuse together slightly distally of the dorsal root ganglion (DRG)/ventral root complex. The complex is a collection of neuronal cell bodies of sensory neurons and axons. The DRG is becoming a therapeutic target for neural interfaces, which primarily aim to selectively interact with the sensory neurons ^58^. Yet, the exploration of cell-body distribution in the DRG has so far remained limited. We found SpIC3D to be capable of capturing details about the cell body distribution within the DRG (**Figure 4E**). We compared SpIC3D to images obtained through histological analysis of the same sample. Visual comparison reveals that both images (i.e., SpIC3D and histology) demonstrate a similar pattern, with a higher cell body density toward the outer circumference on the dorsal side of the DRG. We then quantified cell density over different regions of the DRG (divided into 48 areas) from both histology and MRI images confirming this trend. In both cases, the highest cell body density was observed in the outer ring, toward the dorso-medial side, with a significant difference between the middle and outer rings. Finally, medial and lateral cell body densities were comparable.

In summary, SpIC3D is the first method able to capture critical aspects of the spinal roots’ organization, such as DREZ and AREZ, and to map DRG cell density in anatomically complex scenarios, such as the sacral plexus.

### SpIC3D allows automatic multi-tissue segmentation and 3D reconstruction of the spine

Image segmentation enables extraction of specific tissue features for further quantitative analysis for pathology diagnostics, structural characterization, and interspecies comparisons, as well as 3D anatomical reconstructions for surgical planning and computational modeling. For that purpose, we used a modified HarDNet (**Figure 5A**, detailed in Fasse et al., 2024 ^61^) trained on a small set (< 200 slices, i.e., < 25% of the sample) of manually segmented slices to automatically and simultaneously segment 6 spinal tissues: gray matter, white matter, dorsal roots and rootlets, ventral roots and rootlets, cerebral spinal fluid, and epidural fat. Because of the high bone contrast of the micro-CT images, vertebrae structures were simply segmented by thresholding pixel intensity.

**Figure 5.**
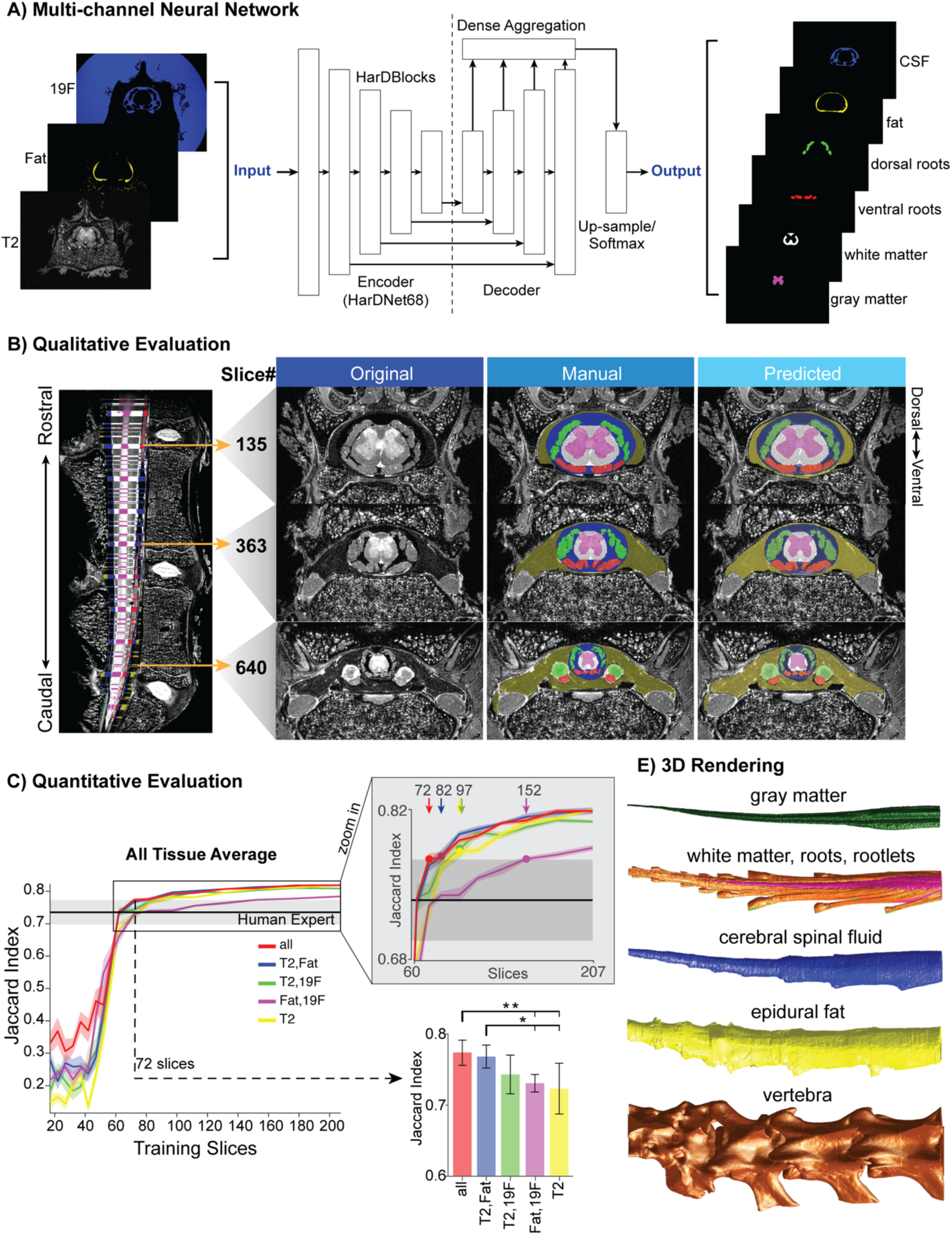
Automatic segmentation algorithm architecture and evaluations. **(A)** Neural network architecture for automatic segmentation, with T2-weighted, fat enhanced, and 19F fluorine MRIs as input, and 6 tissue masks as output: CSF, i.e., cerebrospinal fluid, fat, dorsal roots, ventral roots, white matter, gray matter. **(B)** Qualitative evaluation of segmentation results. *Left*: Mid-sagittal view of T2-weighted image of C01-LS; colored lines indicate image slices that are manually segmented across the spine (207 slices in total) for training and validation of the neural network. *Right*: axial views of 3 selected rostral to caudal slices, comparing the original image, manual segmentation, and automatic segmentation. **(C)** Quantitative evaluation of segmentation results using the Jaccard Indices of the 6 tissue masks (solid line: mean across tissues, shaded area: one standard deviation). Results obtained using different combinations of input channels are compared: all modalities combined (red), T2-weighted and fat enhanced imaging (blue), T2-weighted and 19F (green), fat enhanced and 19F (purple), and T2-weighted imaging alone (yellow) as a function of the number of training slices. A zoomed in plot indicates with dots the number of training slices necessary for each input channel combination to exceed one standard deviation of human expert baseline performance (black line; mean JI of pairwise expert comparison, i.e., a metric for inter-expert variability). Using all imaging modalities (T2, fat, 19F) necessitates the least number of training slices. Bar plot comparing, the average Jaccard Index achieved with 72 training slices for the different input channel combinations (statistical comparison with Kruskal-Wallis test with Bonferroni correction, n=5 for each input condition, significance *: p<0.5, **: p<0.01). **(D)** 3D rendering of 6 tissue geometries obtained using the automatic segmentation neural network, plus the vertebrae.

Qualitative inspection shows good agreement between manual and automatic segmentations (**Figure 5B**). The segmented images capture the rostral to caudal morphological changes in the number and diameter of ventral and dorsal rootlet bundles (**Figure S6A-C**), spinal cord and gray matter size (**Figure S6E, F**), and the rostral to caudal increase in prevalence of epidural fat (**Figure S7**).

To quantify the automatic segmentation quality, we computed the Jaccard Index (JI) and Hausdorff Distance (HD) metrics for the similarity of automatically segmented images and segmentations of the same images by multiple human experts (n=3 experts, 10 sections each), and compared the obtained values with the baseline of inter-operator variability. Specifically, we estimated JI and HD metrics for a batch of neural networks trained with a progressive increased number of training slices (i.e., 20, 41, 62, 82, 103, 124, 144, 165, 186, 207 slices) and using different combinations of input images. The neural networks reached human performance (JI: 0.78, HD: 3.65) after training on 72–152 training slices (depending on the input images; **Figure 4C**). Importantly, using all three MR images (T2w, fat-selective, and 19F for Fomblin) as inputs to the neural network resulted in superior performance compared to that achieved using just one or two. Indeed, the neural networks that used all three MR images required the least number of training slices to achieve human-level performances (72 slices vs. >82) and achieved significantly higher JI and HD values (**Figure 5D**), demonstrating another benefit of SpIC3D.

Finally, three-dimensional renderings of the segmented tissues provide direct visualizations of the sophisticated morphology of the lumbosacral spinal cord, especially regarding the rootlets and epidural fat, captured by the ultra-high-resolution images and segmentations (**Figure 5E**).

In summary, thanks to the ultra-high-resolution imaging data, free of structural distortions, obtained with SpIC3D, we were able to develop the first ever neural network trained to accurately and automatically segment 6 spinal tissues in less than 1 minute, while achieving a comparable accuracy through manual expert segmentation requires 733 +/- 189 hours (see Methods for hardware specifications and human time cost analysis).

### SpIC3D enabled the creation of a highly realistic computational model of the spinal cord

Computational models are a powerful tool for simulating neural responses to clinical and research interventions such as deep brain stimulation and spinal cord stimulation ^62^. For this, a volumetric model with multiple tissue compartments is created and a numerical method, such as the finite element method (FEM), is applied to simulate the electric potential generated within these structures by electrical stimulation. Neuron fiber trajectories and biophysical neural models are then used to estimate neural fiber activation. The anatomical volumes are commonly shaped based on simplified tissue outlines from histology atlases and measures from literature, even though studies have shown that small distortions in tissue compartments such as epidural fat, CSF thickness, and neuron trajectories can give rise to significant differences in simulation predictions ^62–64^. Furthermore, until now neuron fiber trajectories have been mostly manually defined (or based on diffusion simulations ^64^), limiting the accuracy of the simulations.

Here, we exploited 3D volumes (vertebrae, gray matter, white matter, CSF, epidural fat, dorsal roots, ventral roots) and neuron fiber trajectories extracted from SpIC3D to build a highly realistic multi-tissue anatomical computational model for spinal cord stimulation, functionalized with the first ever electrophysiological fiber models based on DTI tracing (**Figure 6A**). To evaluate the benefit of using SpIC3D for the simulation of electromagnetic fields and neuronal recruitment, we compared simulation results against a canonical spinal cord model, in which anatomical tissue contours (**Figure 6A**) and axonal fiber trajectories (**Figure 6C**) were established with approaches commonly adopted in literature, as detailed in the Methods section. When simulating epidural electrical stimulation of the lateral dorsal aspect of the spinal cord, we found significant differences in the voltage distributions and electric field (E-field) strength of the two considered models (**Figure 6B**). The lack of anatomical accuracy of the canonical model led to a substantially lower E-field of up to 96.39% with respect to the highly realistic model obtained with SpIC3D. Importantly, the greatest difference occurred in the vicinity of the dorsal rootlets, which multiple studies have demonstrated to be a crucial stimulation target for spinal cord stimulation ^65,66^. Specifically, the maximum E-field in the dorsal rootlets is 5.8 V/m in the canonical model while 160.6 V/m in the realistic model, with an average of 92.99% difference (**Figure 6B**), which could result in major deviations in simulated neuron activation patterns. Building on methods used in Jantz et al. 2022 ^67^, we then functionalized the two models with realistic (dMRI-derived) and simplified (literature and anatomical measurements, see Methods section) axonal trajectories, and simulated neuronal recruitment (**Figure 6C**). The canonical model, which uses manually defined neural trajectories, predicted significantly higher activation thresholds (by more than one order of magnitude), both for afferent (4.1 vs 0.19 mA median recruitment amplitude for canonical vs. realistic) and efferent fibers (12 vs 0.68 mA median recruitment amplitude for canonical vs. realistic). Experimental data in cats with similar electrode location (L6 vertebral level) and stimulation parameters as used in our models reported nerve recruitment thresholds ranging between 0.12 mA and 0.9 mA, matching those simulated with the realistic model ^67,68^. This suggests that the canonical model strongly overestimates the amount of current necessary to stimulate the fibers (**Figure 6D**). Interestingly, also the activation pattern differed dramatically between the two models: the canonical model predicted a greater separation between the afferent and efferent activation thresholds, suggesting an over-estimated selectivity not supported by the highly realistic model. While it is experimentally challenging to distinguish between afferent and efferent fiber activation, spinal cord stimulation experiments in low-lumbar cord that analyzed stimulation-evoked H-wave (elicited by afferent fibers activation) and M-wave (elicited by efferent fibers activation) responses agree with a predominantly afferent-first activation, but do not suggest a complete separation between afferent and efferent responses ^69,70^. Importantly, these fiber activation pattern differences as well as the underestimation of the extracellular voltage and E-field due to anatomical inaccuracies would result in an incorrect selection of the stimulation parameters. These could all be potential contributing factors to the superior neuromodulation effectivity using parameters optimized with personalized models compared to canonical models in the clinic ^64,71^.

**Figure 6.**
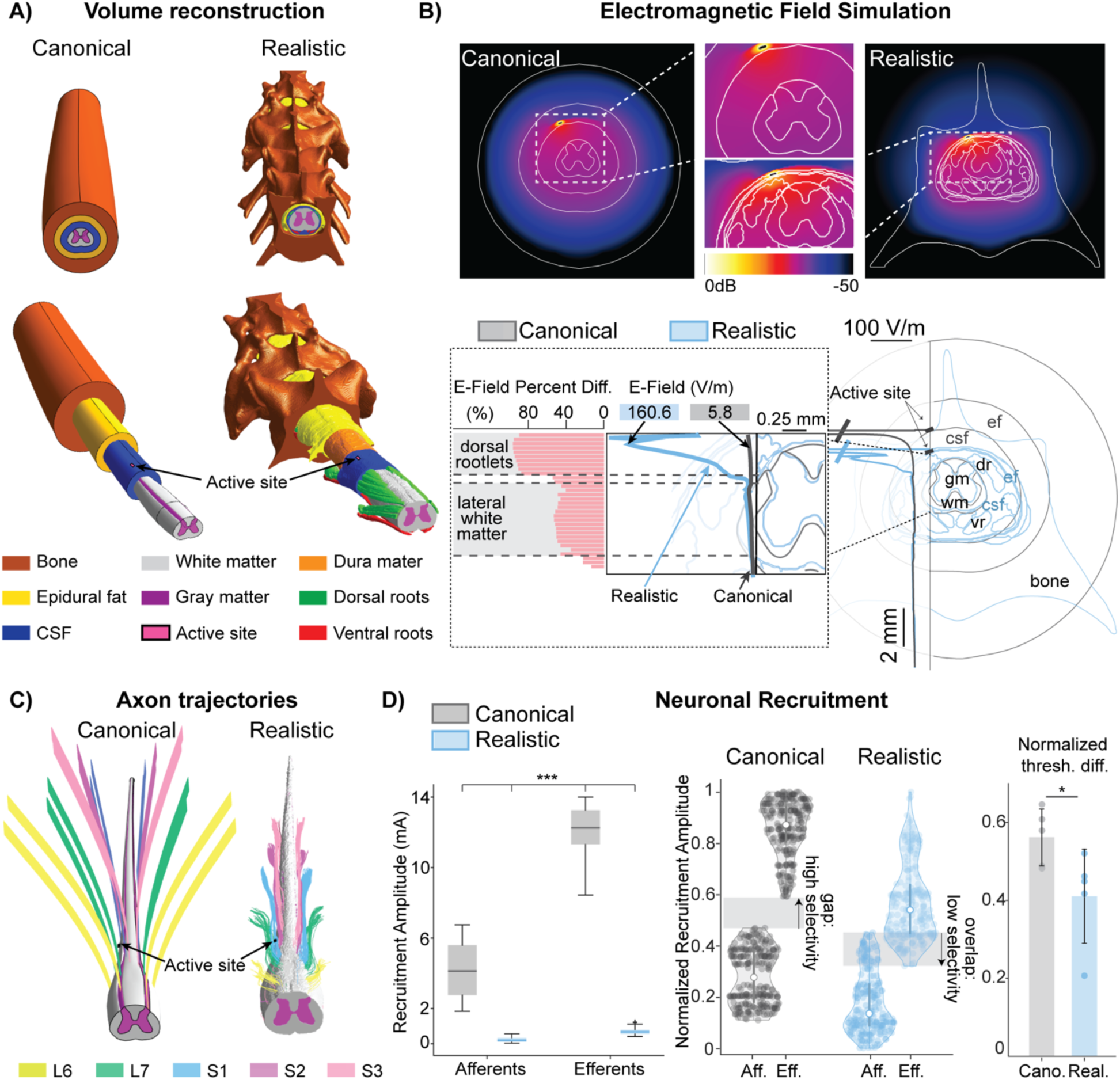
Canonical versus highly realistic lumbosacral spinal cord model of spinal cord stimulation comparisons. **(A)** *Left*: a canonical computational model constructed based on anatomical dimensions from MRI and literature using extrusion, featuring gray matter, white matter, epidural fat, CSF, and bone tissue compartments. *Right*: a highly realistic computational model constructed from high-resolution MRI and CT image segmentations, containing dorsal/ventral roots and dura mater in addition to all tissue compartments in the canonical model. The active sites for spinal cord stimulation (SCS) are placed inside the lateral epidural fat, with one side in contact with either the CSF or dura matter. **(B)** *Top*: Simulated electromagnetic field maps on the level of the active site (axial plane) for both the canonical and the highly realistic model, with consistent unitary voltage input from the electrode. Insets highlight the faster potential drop-off (higher E-field strength) close to the active site in the canonical model. *Bottom*: plot of simulated E-fields for canonical (gray) and realistic (blue) models along the dorso-ventral direction marked by a vertical line. Inset shows the percent E-field difference (pink plot) between the two models at corresponding locations on the spine. The greatest difference occurs where the dorsal rootlets reside. **(C)** Axon trajectories used for neuron fiber recruitment simulations. *Left*: dorsal and ventral root axon trajectories for the L6-S3 roots of the canonical model, modeled with splines based on dissection-measured root angle orientations. *Right*: realistic axon trajectories of the highly realistic model, extracted from DTI tractography for L6-S3. **(D)** Afferent and efferent neuron recruitment comparison between the two models (n=500 neurons each, diameter of 12 µm, evenly split into afferents and efferents). *Left*: Box plot of absolute neuron recruitment amplitudes (mA) obtained from the canonical model, which are significantly higher than those from the realistic model for both afferents and efferents. (Kruskal-Wallis test with Bonferroni correction used for statistical comparison, *: p<0.05, **: p<0.01, ***: p<0.001) *Middle*: Violin plot of neuron recruitment amplitudes normalized to its highest value for each model to compare afferent/efferent selectivity by SCS. *Right*: Bar plot comparing afferent and efferent normalized activation threshold differences for the two models for each root level (L6-S3, n=5). The canonical model has a significantly higher activation threshold difference than the realistic model (Mann-Whitney U-Test used for statistical comparison, *: p<0.05). For all boxplots, the whiskers extend to the maximum and minimum, excluding outliers. Central line, top, and bottom of the box represent median, 75th percentile, and 25th percentile, respectively.

Overall, these results suggest that a highly realistic computational model of the spinal cord obtained with SpIC3D could significantly improve the accuracy of simulated electromagnetic fields and neuron recruitment patterns compared to commonly used modeling approaches, supporting the development of spinal cord stimulation protocols with superior safety and effectivity.

### SpIC3D generalizes to the cervical and thoracic spine

There are significant morphological differences between the cervical, thoracic, and lumbosacral spine and spinal cord within the same species, which so far have hindered the generalization of imaging protocols across segments ^72^. For instance, in addition to differences in root exit angle and direction of the roots, the size of the cord and the gray/white matter ratio significantly diminishes rostro-caudally (**Figure 1B**). It was, therefore, critical to validate the applicability of SpIC3D across the entire spine.

For this, we imaged a cervical and a thoracic spine segment of an adult cat using SpIC3D (imaging field of views shown in **Figure 1B**). With the same in-plane resolution for CT (100 µm), MRI (50-100 µm) and dMRI (100 µm), and same diffusion directions for dMRI (19 directions, b-values 1000 s/mm² and 2000 s/mm²) as used in the lumbosacral tissue, we successfully applied SpIC3D to both cervical and thoracic spine tissue (**Figure 7**). The 3D volume reconstruction of the images clearly showed that, similar to the lumbosacral spine, SpIC3D accurately captured the fine details of the cervical and thoracic anatomical structures. More specifically, the images revealed thicker lateral and dorsal epidural fat in the thoracic spine as compared to the cervical and lumbosacral spine (**Figure S7A-B**), as expected ^73^. Additionally, we detected expected differences in gray matter sizes with respect to the spinal cord (gray matter proportion) between cervical, thoracic, and lumbosacral spine. Specifically, the thoracic cord had the smallest and the sacral cord had the largest gray matter proportion (**Figure 7C**), congruent with literature ^74^. Importantly, both volume reconstruction and fiber tractography of the spinal roots were consistent with observations from dissection and histology ^75–78^. Indeed, cervical roots were larger in diameter (**Figure 7C**), and they exited the spine at nearly the same rostro-caudal location as their DREZ (**Figure 7A**). Thoracic roots, instead, were substantially smaller in diameter (**Figure 7C**), had more intersegmental space, and exited the spine either at or a little below their DREZ (**Figure 7B**). Therefore, as expected, the root angles from the rostro-caudal axis for both cervical and thoracic roots were much larger than those of the lumbar and sacral roots ^3^ (**Figure 7C**).

**Figure 7.**
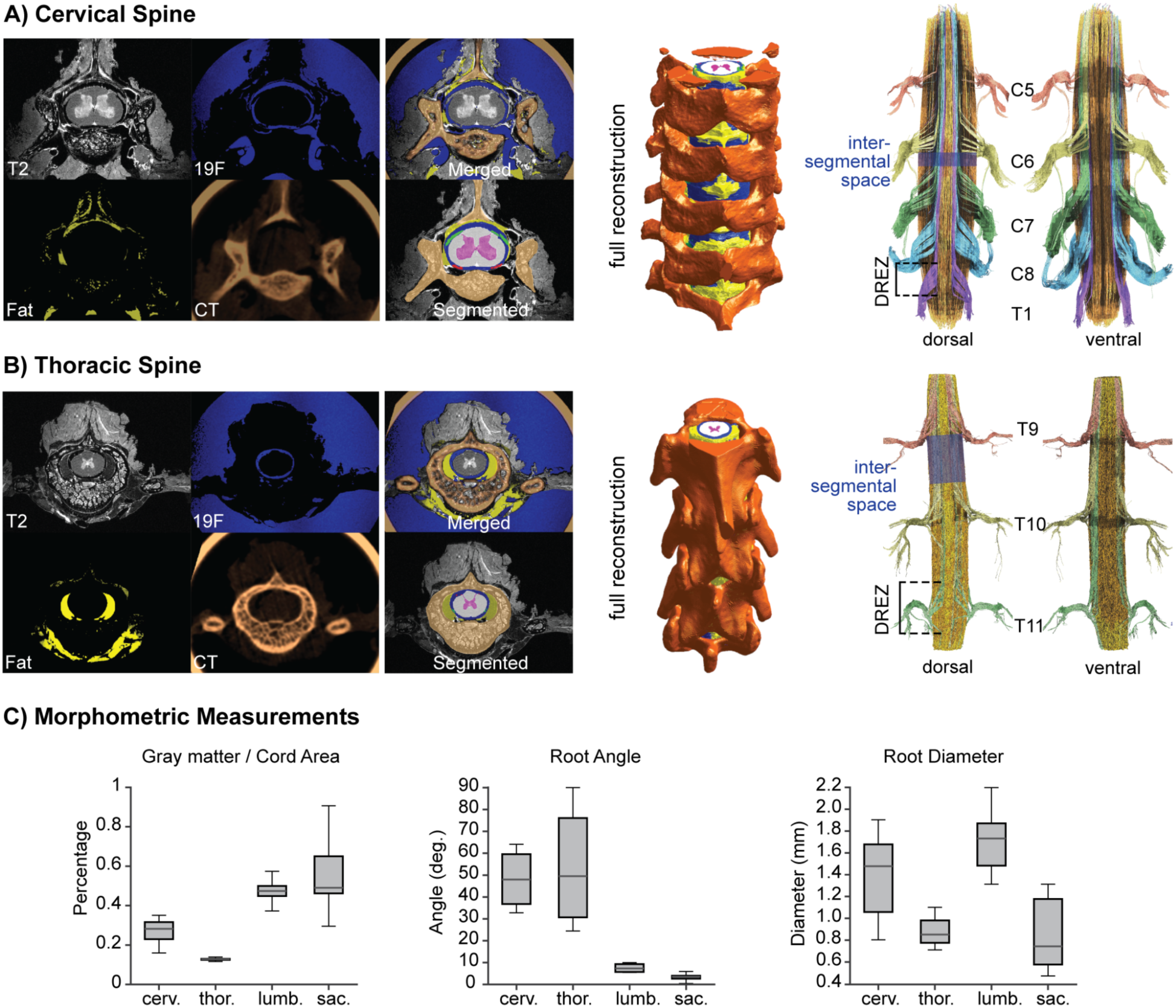
SpIC3D of the cervical and thoracic spinal segments of a cat. (A), **(B)** *Left*: axial view of C05 cervical and thoracic spine MRI and CT images, as well as the merged image of all sequences, and corresponding segmented tissue layers (pink: gray matter, white: white matter, green: dorsal roots, red: ventral roots, dark blue: CSF, yellow: epidural fat, copper: bone). *Middle*: full 3D reconstructions of the spine segments (C4-T1, T9-T11) with all segmented tissue compartments. *Right*: dorsal and ventral views of DTI tractography performed from ROIs around each spinal root. Fibers are grouped by color corresponding to each root. DREZ and intersegmental space can be visualized from the fiber reconstructions. **(C)** Boxplots showing MRI measured quantitative comparisons across cervical (left and right C5-C8; n=8), thoracic (left and right T1, T9-T11; n=8), lumbar (left and right L5-L7, from samples C01, C03, C06; n=6 per sample) and sacral (left and right S1-S3, from samples C01, C03, C06; n=4,6,6 respectively) segments. From left to right, gray matter proportion in the spinal cord, root angle, and root diameter. For all boxplots, the whiskers extend to the maximum and minimum. Central line, top, and bottom of the box represent median, 75th percentile, and 25th percentile, respectively.

In summary, SpIC3D is the first ultra-high-resolution volumetric multistructural imaging protocol that generalizes across spinal segments without adjustment in the acquisition protocols.

### SpIC3D generalizes to human samples

Finally, to demonstrate the scalability and translational potential of SpIC3D to humans, we applied the pipeline to a human cadaver thoracic spine (**Figure 8A**). The human spine is not a homogeneously scaled up version of the cat spine. Indeed, while the human spinal cord is approximately 1.25 times the diameter of the cat spinal cord, the lumbar enlargement is approximately 2 times the length and the width of the spinal canal can be more than 3 times that one of the cat ^79^. Additionally, the morphological differences in the spinal canal between cats and humans are reflected in the size and shape of the fat and CSF. We implemented SpIC3D in a 9.4T scanner with a 12 cm gradient set to accommodate the larger vertebral size. These dimensional differences created challenges for imaging, yet SpIC3D was implementable in a 9.4T Bruker scanner and it could accurately capture the organization of the different spinal structures allowing a volumetric reconstruction of the human spine (**Figure 8C-D**). The organization of the roots is also different between the two species. Since the conus medullaris is at different vertebral levels in cats and humans, i.e., S3 and T12/L1, respectively, rootlets are more tightly packed in the human spine and travel parallel to each other for a much longer distance before exiting the spinal canal. Regardless, the resolution and optimization of the 19-direction dMRI in SpIC3D generalized across species and morphological differences. Indeed, also in the human sample, we could visualize individual fiber bundles, trace them back to label the different roots, and quantify the DREZ, CSF and epidural fat distribution, as well as gray matter and white matter areas (**Figure 8B, E, Supplementary Table 2**). No previous MRI methods in human subjects were able to differentiate roots at the thoracic-lumbar level where multiple tiny roots and rootlets with the same orientation are present. Importantly, we were able to quantify the diameter of the different roots (**Supplementary Table 2**) which match with various previous dissection reports from literature ^3,75,76^.

**Figure 8.**
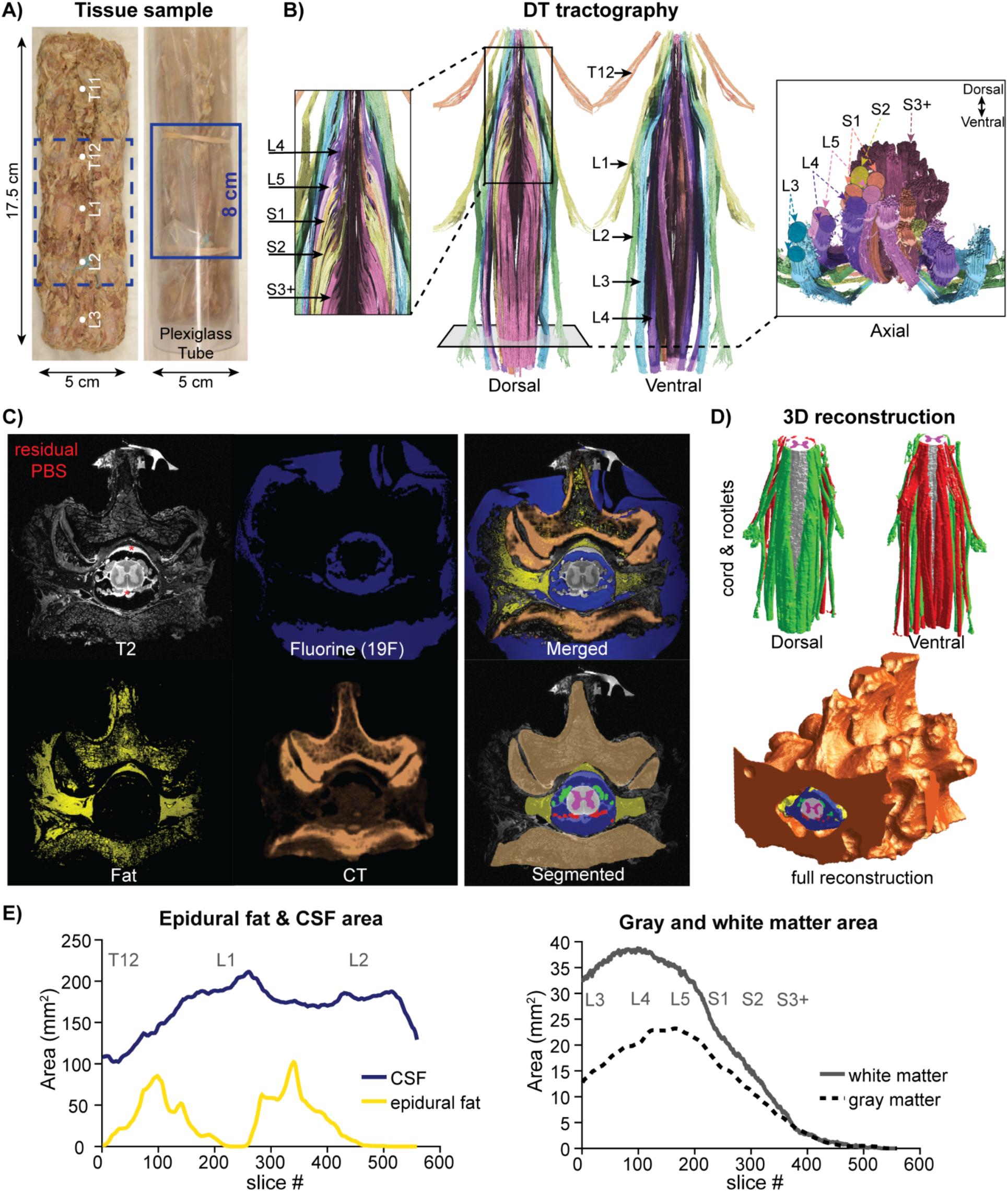
SpIC3D of a human cadaver thoraco-lumbar vertebral segment (lumbosacral cord segment). **(A)** Thoraco-lumbar spine sample (T11-L3) extracted from a human cadaver (H01), with T11 and L2 vertebrae marked with sutures. The sample is also shown packed in a vacuum bag filled with Fomblin, placed in a 5 cm plexiglass tube for imaging. The 8 cm imaging field of view (FOV) is centered at the L1 vertebra, shown by the blue solid or dashed box. **(B)** Dorsal and ventral views of DTI tractography performed from ROIs drawn around each spinal root, and around the spinal cord. Fibers are grouped by color corresponding to each root. *Left inset*: DREZ visualized for L4-S3+ levels with relatively clear distinction between levels but also some overlap. *Right inset*: axial view from the caudal side of the spinal canal, with approximately groups of 2 root bundles (dorsal root and ventral root) on each side of the cord for each of L3-S1 root levels. Those for S2 and S3+ levels are still in rootlet form, presenting multiple small diameter fiber bundles per root level. **(C)** Axial view of H01 spine MRI and CT images, as well as the merged image of all sequences, and corresponding segmented tissue layers (pink: gray matter, white: white matter, green: dorsal roots, red: ventral roots, dark blue: CSF, yellow: epidural fat, copper: bone). **(D)** 3D reconstructions of the spine segment (T12-L2) displaying the spinal cord with dorsal and ventral rootlets (*top*), and full reconstruction with all segmented tissue compartments (*bottom*). **(E)** *Left:* Plot showing CSF and fat areas across the imaged length of the H01 spine. T12, L1, L2 labels correspond to the vertebral levels. *Right:* Plot of gray and white matter areas across the imaged length of the H01 spine. L3, L4, L5, S1, S2, S3+ labels correspond to spinal cord levels.

In summary, SpIC3D is the first ultra-high-resolution volumetric multistructural imaging protocol that generalizes to the human spine allowing fine reconstruction of thoracic-lumbar spinal rootlets.

## DISCUSSION

Imaging the structural organization of the central nervous system at a resolution that is sufficient to distinguish cell distributions and white matter fiber bundles is quickly becoming a priority in neuroscience research and clinical practice. While an abundance of effort has been put into optimizing brain imaging towards this goal, advancements in spinal cord imaging are still limited. Here, we designed a postmortem in-situ high resolution volumetric and multistructural imaging pipeline (SpIC3D) that enabled the most complete characterization and quantification of spinal cord samples to date. In particular, we propose a novel application of a 19F MRI for Fomblin to image a void space. This allowed first-ever characterization of the CSF for ex-vivo imaging while reducing noise in other MRI sequences. Additionally, spinal quantifications as those reported here were previously achievable only through multiple studies involving microdissections, histology, and imaging methods combined. With these features, SpIC3D opens new perspectives for spinal anatomical characterization. We discuss our findings with an emphasis on its applications in neuroscience research, clinical practice, and its translation towards in-vivo imaging.

### Neuroscience Research Applications

The SpIC3D pipeline can be a valuable tool for spinal cord and nerve research applications in multiple aspects. Not only were we able to visualize and quantify the DREZ and AREZ with SpIC3D, but we were also able to see the major afferent trajectories into the gray matter and their rostro-caudal span within the gray matter with respect to the DREZs and AREZs. This brings interesting new perspectives on the definition of spinal segment levels. Indeed, there are two widely used methods to define spinal segment levels: 1) organization of the spinal motoneuron pools ^80^, that is more commonly used by neuroscientists, and 2) insertion border of (dorsal) spinal nerve roots into the spinal cord ^81^, which is widely used in neurosurgical practice; however, it is unclear how well these two definitions correspond with each other with critical implications for surgical procedures and neuromodulation approaches. From our SpIC3D fiber reconstructions, we found that the span of the afferents within spinal gray matter, which could correspond to primary afferent arborization around the motoneuron pools, is wider and more rostral than the DREZ. This finding is in agreement with retrograde labeled afferent trajectories for cat lower limb muscles reported by Ishizuka et al. 1979 ^59^. This location mismatch between the afferent fiber entry zones and their corresponding effector motoneurons implies a possible functional mismatch between the two prevailing definitions of spinal segment levels. It also highlights the need to better quantify this mismatch at a population level, which could be achieved with the assistance of SpIC3D.

Second, the SpIC3D pipeline enabled us to successfully quantify cell distribution within multiple levels of the DRG as effectively as histological analysis. Importantly, we tested this critical ability of SpIC3D to resolve neural elements like cell bodies in a relatively simple structure, as the DRG only contains cell bodies and axons. However, we do expect that SpIC3D could generalize to more complex structures. Additionally, animal models are frequently used to improve DRG based treatment and neuromodulation therapies; however, the similarity and differences of cell distribution between these models and humans are still unclear, hindering translation and optimization of these therapies. The SpIC3D pipeline could empower rapid and distortion free characterization and comparison of DRGs across multiple species resolving this controversy. Indeed, SpIC3D showed high generalizability across multiple samples, multiple spinal segments (cervical, thoracic, lumbar), and multiple species (cat and human) despite their intrinsic physical variations. Importantly in all the samples, we were able to perform morphological quantifications repeatedly, and more comprehensively than what can be measured through dissection (**Supplementary Table 3**). As a result, we were able to obtain one of the most comprehensive postmortem spinal cord datasets available. In this regard, SpIC3D provides all features required to build personalized finite element models of the spinal cord, which could be used to simulate electrical responses as shown here, or mechanical responses to assess cause of injury or device safety, reducing or replacing the need for animal use.

Finally, the SpIC3D pipeline will facilitate the maximization of tissue use for multiple purposes, in line with the “three Rs” principles in research: Replacement, Reduction, and Refinement ^82,83^. Specifically, in this study we first performed in-situ imaging of the spine samples, then, we performed dissection measurements, followed by histology processing and analysis of the DRG cell distributions. During this process, we did not notice any damage or distortion to the tissue that affected the quality of the downstream analysis. While in our study we performed these steps mainly for the direct validation of our imaging results, others could use a similar approach (i.e., first imaging and then tissue processing) for complementary analysis such as immunohistology of the same samples, which would effectively reduce the total number of animal samples required. In the same direction, we made all imaging datasets obtained in this study openly available for the neuroscience community.

### Clinical Applications and In-Vivo Translation

While interest in spinal cord imaging is growing rapidly, there has been a lack of standardized and comprehensive analysis framework, limiting progress in its clinical potentials. This idea was at the base of the creation of the spinal cord toolbox (SCT), i.e., a comprehensive, free and open-source set of command-line tools dedicated to the processing and analysis of spinal cord MRI data ^84^. SpIC3D provides an excellent tool to provide a more accurate and comprehensive atlas for the SCT. For example, people in the spinal functional MRI (fMRI) community recognized recently that spinal roots and rootlets are critical in determining correct spinal segments rather than relying on vertebral levels ^3,81,85^. The SPIC3D will allow us to revisit the segmental organization of the spinal cord. Indeed, high-resolution MRI and DTI provide visualization of both intraspinal gray matter organization and the extraspinal nerve roots and rootlet entry zones, and with enough resolution even intra gray matter fiber trajectories. This information could be inserted into SCT improving its pipeline to guide more precise and functionally relevant spinal cord fMRI analysis ^81^.

Additionally, SpIC3D could have strong implications for neurosurgical procedures such as nucleus caudalis DREZ lesioning ^86^, or for postmortem pathologic assessment of the spinal cord at autopsy, which relies on anatomical post-mortem atlas. Indeed, the automated segmentation and morphometric analysis achievable with SpIC3D yielded results similar to previous experiments based on manual measurement of individual tissue slices. Finally, ablation of spinal tissue to alleviate neuropathic pain can also be performed at the level of the DRG. The analysis of cell distribution within the DRG obtained here thanks to SpIC3D could help increase the specificity for DRG stimulation or ablation ^58^.

Finally, generalization of SpIC3D to in-vivo imaging could further and improve neurosurgical planning. For instance, fast and automatic segmentation of tissues via a deep neural network as the one developed here, and with the assistance of fiber tractography, thanks to SpIC3D, could help in precisely detecting boundaries of intramedullary and spinal Schwannoma tumors and improve resection planning to avoid damaging critical white matter tracts ^87^. Similarly, high resolution visualization of the rootlets’ organization and particularly of the DREZ could allow more precise identification of damaged nerve roots for ablation to treat neuropathic pain and ameliorate placement of SCS electrodes. Indeed, nowadays the implantation is guided by counting the vertebra ^2,48^. However, neuroanatomical organization of root and rootlet fibers, i.e. the primary target of SCS, within the vertebrae varies across individuals ^3,64,88–90^. Therefore, personalized implantation based on fibers organization could improve clinical outcome of SCS to treat pain and restore sensorimotor and autonomic functions in spinal cord injury patients ^62,71^. Importantly, we demonstrated that SpIC3D and our neural network generalizes to other spinal segments ^61^ and species creating the foundation for this future development. Furthermore, we believe that the optimization of gradient directions and b-values performed here would generalize to in-vivo tissues and could therefore be used for these purposes.

In conclusion here we presented a spinal cord in-situ high-definition volumetric multistructural imaging method that allowed quantification of neuronal cell density in dorsal root ganglia, multi-segment identification of individual rootlets and roots, and volumetric reconstruction of multiple spinal structures at unprecedented resolution. This highly versatile tool provides a basis for accelerated spinal pathology characterization and personalized spinal cord stimulation treatments.

## METHOD DETAILS

### Cat Postmortem Spine Tissue

#### Sample Extraction

We used six perfused and formalin preserved cats (Ward’s Science 470154-114/470154-122 Uninjected Skinned Cats), which we named C01-C06 in this manuscript. To extract the spine samples, we first exposed the cat spinous processes with a dorsal medial incision along the spinous process to expose the vertebral column, then located the vertebral levels of interest (cervical enlargement: C3-T2, thoracic: T8-T12, lumbosacral: L3-S3) by counting the spinous processes. We placed sutures on either T8 and T13 spinous processes for thoracic samples, and L5 and L7 spinous processes for lumbosacral samples to assist with centering the field of view.

For samples C01 and C03-C06, we carefully removed the muscles and connective tissue surrounding the vertebrae and cut any nerve roots extending out of the vertebral column with a scalpel to prevent disturbance to the roots within the vertebral column. Finally, we transected the spinal cord and vertebral column above and below the segments of interest to detach the sample, keeping the bone intact. After extraction, we checked that the sample was slightly less than 3 cm in diameter and fit snugly inside a 30 mm sample glass tube (Bruker) that fit a 30 mm RF probe for imaging. Most samples fit in the tube without alterations to the bone, although for some lumbosacral samples, we had to trim down 2-5 mm from the transverse vertebral processes. For sample C02, we dissected a longer and wider piece of tissue (**Figure S5B**), from L1 to the base of the tail, and laterally out to include the sciatic and pudendal nerve branches. We identified the pudendal nerve as a small ∼2mm nerve that branches medially from the sciatic nerve that innervates the pelvic floor muscles, and we marked it with sutures.

#### CT Image Acquisition

To capture the 3D structure of the vertebral column, we obtained computed tomography (CT) images of the spine samples using a small animal CT scanner (Si78; Bruker BioSpin GmbH, Ettlingen, Germany) equipped with the software package ParaVision-360 (version 3.2; Bruker BioSpin Corp, Billerica, MA, RRID:SCR_001964, http://www.bruker.com/service/support-upgrades/software-downloads/mri.html). To ensure accurate registration with the MRI, we kept the specimens inside the glass sample tubes for the CT to avoid any undesired vertebral curvature. We used a “Low Dose” 1 mm aluminum filter and a “step and shoot” method (0.6-degree gantry step), and the images (100 μm isotropic, FOV: 79.6 x 79.6 x 154.7 mm) were reconstructed using a filtered back projection algorithm.

#### Tissue Preparation for MRI

Prior to MRI, the samples were soaked in a Gadolinium contrast agent solution to reduce the tissue T1 relaxation and afford rapid 3D imaging acquisition. Samples were placed in a 1x phosphate-buffered saline (PBS) (Sigma Aldrich, St Louis, MO, USA) solution doped with 0.2% Gadavist (Bayer Health, Leverkusen, Germany) for two weeks. The samples were then placed in 0.2% Gadavist in dH2O to partially recover the T2 reduction to ensure good contrast ^50^. After 24 hrs, the aqueous solution surrounding the samples was replaced with Fomblin (Solvay Specialty Polymers, Bollate, Italy) for magnetic susceptibility matching with a proton-free background. The samples were degassed under gentile vacuum prior to imaging. Fomblin has a density of 1.8 g/mL while PBS solution has a density of 1.005 g/mL ^91^ allowing Fomblin to sink to the bottom and easily replace any remaining PBS solution and air to avoid artifacts.

#### MR Image Acquisition

High-resolution images were collected using a 11.7 Tesla/89 mm vertical-bore actively-shielded Bruker AVANCE AV3 HD microimaging system, equipped with either a Micro2.5 gradient set and 30 mm QTR birdcage resonator or Mini0.75 gradient set and 40 mm QTR birdcage coil. The 40mm resonator and MINIWB57 animal handling system allowed for translation of the spine in the RF coil to increase the FOV by sliding the coil to acquire two overlapping 4.4 cm imaging sets, and tiling two sets of images together (samples C03-LS, **Figure 2**). For each sample, four 3D data sets were collected. A 3D T2 RARE (rapid acquisition with relaxation enhancement) for tissue contrast, a fat-selective RARE with a narrow RF pulses (1 kHz BW) centered at −3.5 ppm, a 19F RARE (with the carried frequency at −81.7 ppm) for detection of the Fomblin in the CSF void, and a DTI data set with one or two shells where the diffusion directions were selected using the Emmanuel Caruyer online q-space sampling protocol application. The application uses a multi-shell generalization of electrostatic repulsion to sample uniformly distributed directions in the q-space ^53^. Details of acquisition parameters are found in supplementary material and **Supplementary Table 1.** In general, the FOV was 40 x 19.2 x 25.6 mm and T2w was collected at 50 µm isotropic resolution or 50 µm in-plane and 100 µm thick, Fat-selective MRI at 50 or 100 µm resolution, and DTI at 100 µm, 150 µm or 200 µm resolutions. For 19F MRI, we changed to a two-channel 1H/19F resonator and the FOV was increased to 44 x 34 x 34 mm to avoid fold-over of the Fomblin signal surrounding the spine sample. 3D 19F images were collected at 100 µm resolution and an additional 1H RARE was collected to assist with co-registration. Due to the larger size of the C02-LS sample, it was scanned instead in the 9.4 Tesla 30-cm Bruker Biospec AV3 spectrometer equipped with an BGA-12S2 HP gradient insert and a 72 mm quadrature RF resonator.

### Human Postmortem Spine Tissue

We obtained one thoraco-lumbar spine sample from a formalin fixed (unknown duration) cadaveric human (H01-LS), without any known neurological diseases, from the University of Pittsburgh School of Medicine anatomy lab (protocol approved by CORID, ID: 1070). To extract the sample, we placed the cadaveric specimen in a prone position and identified the desired vertebral segments (T12-L2) hosting approximately the L3-S3 spinal segments by counting the spinous processes. We started the dissection with a dorsal incision along the side of the spinous process, and carefully dissected laterally to expose the vertebral column. Then, we cautiously transected the spinal column at least one vertebra above and below the desired segments (T11-L3). Thereafter, we cut the lateral muscles and connective tissue to detach the vertebral segments of interest, with bone intact. Once the segments were detached from the rest of the cadaveric human, we trimmed the muscles until the sample was about 5 cm in diameter, and marked the T11 and L2 spinous processes with different colored sutures as reference for the rostro-caudal orientation.

We acquired the CT images the same way as for the cat samples. Since the cadaver spine sample is larger in size, we kept it in the 1x PBS + 0.2% Gadavist solution for three weeks before scanning to ensure sufficient rehydration and gadolinium penetration. Due to its larger size, the sample was thermally sealed in a polyethylene bag filled with Fomblin and placed in a 5 cm plexiglass tube for imaging. The MRIs of the sample were acquired in the 9.4 Tesla 30-cm Bruker Biospec AV3 spectrometer equipped with an BGA-12S2 HP gradient insert and a 72 mm quadrature RF resonator. Just like for the cat spine samples, we acquired a 3D T2 RARE, a fat-selective RARE (−3.5 ppm), a 19F RARE, and a set of DTI images. 19F MRI used a dual-tuned 1H/19F resonator with the carrier frequency at −81.7 ppm. Details of acquisition parameters are documented in the **Supplementary Table 1.**

### T1 and T2 Relaxation Time Monitoring

For the C03 lumbosacral, and the C05 lumbosacral, thoracic, and cervical samples, the tissue T1 and T2 relaxation times were monitored during the gadolinium preparation period. For C01 lumbosacral and H01 samples, the initial pre-gadolinium and final post-gadolinium treatment relaxations times were recorded without intermediate monitoring. We measured tissue T1 and T2 relaxation time of C03 and C01 lumbosacral samples using a 11.7 Tesla/89 mm vertical-bore actively shielded Bruker AVANCE AV3 HD microimaging system, equipped with a Mini0.75 gradient set and 40 mm QTR birdcage coil. For C05 and H01 samples, we used a 9.4 Tesla 30-cm Bruker Biospec AV3 spectrometer equipped with an BGA-12S2 HP gradient insert and an 86 mm quadrature RF resonator. We stacked all C05 samples and imaged with a 60 x 60 mm FOV and 255 x 256 matrix, 2 mm slice thickness and 11 slices. In general, T1 was measured with a variable TR spin-echo sequence, with 6 images sampling TR between 50 and 5000 ms, and T2 was measured with a multi-echo spin-echo with a TR of 4000 ms and TE 8 with 30 echo images paced at 8 ms. T1 and T2 were measured separately for gray matter and white matter, then they were fitted to a mono-exponential recovery or decay, respectively.

### Image Segmentation

#### Image Preprocessing and Manual Segmentations

Segmentations of spinal tissue layers were performed using the FMRIB Software Library ^92^ (FSL, University of Oxford). Before segmentation, we first co-registered and resampled the Fat-Enhanced, 19 Fluorine, and CT imaging sequences to the 50 µm in-plane T2-weighted scans using the Nudge tool in FSLeyes. The resulting images were then layered for concurrent visualization to facilitate segmentation. Three trained specialists (LL, UA, CR) familiar with the spine anatomy performed manual segmentations for separate spine samples based on the T2-weighted, Fat-selective, and 19 Fluorine MRIs. Due to the ultra-high spatial resolution, each image stack contained 600-800 axial slices, thus manual segmentation was performed only on select evenly spaced slices (ranging between 40-207 slices) representative of spinal morphology across the rostro-caudal spine, which served as ground truth for the automatic segmentation algorithm^61^. Six tissue masks of the spine, including 1) gray matter, 2) white matter, 3) cerebrospinal fluid (CSF), 4) epidural fat, 5) dorsal roots, and 6) ventral roots were segmented, without tissue overlaps, using the Edit tool in FSLeyes. The coregistered CT images provided excellent bone tissue contrast, thus, we segmented the full vertebrae volume using the free open-source image computing software, 3D Slicer ^93^ (https://www.slicer.org/), through simple intensity thresholding.

#### Manual Segmentation Time Cost Analysis

With higher-resolution imaging comes a linear growth in the number of image slices for a tissue, and thus image slices to be segmented to reconstruct the tissue for analysis. Detailed data on time spent performing segmentations was collected from the three segmentation specialists. The segmentations were performed on the image set C01-LS, which consists of an anatomical T2-weighted, a Fat-selective, and a 19F Fomblin image of L5-S3 of the cat spinal cord, totaling 800 slices. The participants used these images to identify white matter, gray matter, CSF, epidural fat, dorsal rootlets, ventral rootlets, dorsal roots, and ventral roots. Participants were asked to collect the time to segment the 8 tissues for 5 to 8 set of 10 slices. In order to account for learning effects, we then considered only the time of the last set and we multiplied it by 80 to estimate the total time to manually segment the full 800 slice spine volume. For our trained segmentation experts, we calculated that it would take them 800 hrs, 880 hrs, and 520 hrs respectively, averaging to 733 +/- 189 hrs.

#### Segmentation Accuracy Assessment Metrics

To assess the accuracy of the segmentation we used the Jaccard index J, which is an overlap area-based metric. It is defined as 𝐽=|𝐴∩𝐵|/|𝐴∪𝐵|, where A and B are two segmented masks being compared. A higher Jaccard index signifies a higher overlap, and thus a greater correspondence between ground truth and prediction. The second accuracy metric employed is the Hausdorff distance H, which is a spatial distance-based metric. It is defined as the maximal (pixel) separation of the boundaries of A and B. A smaller H value signifies a better boundary fidelity. Deviations in boundary locations may ultimately have important impacts on downstream analysis such as nerve recruitment predictions, and the Jaccard index is less sensitive to such deviations (especially when the segmentation has a high volume to boundary ratio).

#### Baseline Human Expert Performance

To facilitate interpretation of the accuracy metric results, we collected inter-operator segmentation variability as a reference benchmark for ideal performance. All three segmentation specialists were asked to segment the tissue masks on the same ten slices of the image set C01-LS. The Jaccard index and the Hausdorff distance of their segmentations were calculated pairwise, and we computed the average scores to provide the human performance baseline.

#### Neural Network and Automatic Segmentations

Automatic segmentation was achieved through a novel Convolutional Neural Network (CNN) based image segmentation pipeline ^61^. This pipeline involves bias field correction (N4ITK), volumetric masking, and intensity normalization (Nyul) for preprocessing. Then, the preprocessed images and manually segmented axial slices were used as the input and ground truth respectively for training the modified HarDNet MSEG neural network which uses a Dice Loss function (see Fasse et al. 2024 ^61^ for more details). We kept 20 slices of manual segmentation aside to validate the training results. This process takes an average of 14 minutes in a GPU-based, CUDA-enabled python environment on a machine equipped with 32-core AMD Threadripper 3970X with 256GB of DDR4 RAM and two Nvidia GigaByte RTX 3090 24GB GPUs with NVLink. Finally, the trained neural network predictions were compared to the 20 validation slices to report the Jaccard score and Hausdorff distances. In this study, we used 207 manual segmentation slices from C01-LS and 176 slices from C03-LS to train the neural network, which was used to predict all 6 tissue segmentations for C01-LS (800 slices), C03-LS (770 slices), C06-LS (796 slices), as well as C05-Ce (800 slices) and C05-Th (800 slices) with transfer learning (see Fasse et al. 2024 ^61^ for more details). For the cadaver sample H01 (656 slices), we retrained the network (due to image dimension differences from the cat images) with 44 manual segmentation slices and used that to predict all 6 tissue segmentations in all 656 slices.

#### Neural Network Input Channel Combination Analysis

All possible input channel combinations from C01-LS were considered in this analysis to determine which input channels are useful for the neural network to perform good segmentations. This consisted of 1) all 3 MRI inputs, 2) T2w and fat-selective images, 3) T2w and 19F Fomblin images, 4) fat-selective and 19F Fomblin images, 5) T2w image only, 6) fat-selective image only, and 7) 19F Fomblin image only. Then, for each condition, 17 different numbers of training slices were used to train the neural network (n=17, 22, 27, 32, 37, 42, 47, 52, 62, 72, 82, 97, 112, 132, 152, 177, 207). The set of training slices were randomly selected from the total 207 manually segmented slices. The neural network was trained for 30 epochs with 3 augmentations per slice, and repeated 5 times for each condition, i.e., each number of training slices. The Jaccard score was calculated for each tissue layer for each training, and the mean and standard deviation of the scores were exported in an excel file and visualized in MATLAB (version R2020b).

### Diffusion Tensor Image Analysis

#### DTI Fiber tracking and coordinate extraction

All diffusion tensor reconstruction, fiber tracking, and diffusion metrics extraction were completed using the open-source software DSI Studio (http://dsi-studio.labsolver.org, CUDA SM75 version Apr 7, 2022). Signal intensity threshold was set to 50 and a DTI Gaussian model was used for tensor reconstruction. The b-table was checked by an automatic quality control routine to ensure its accuracy ^94^. When running the deterministic tractography ^95^, the anisotropy threshold, angular threshold, and step size were set to default values. A minimum tract length was set to 10 mm and maximum tract length was set to 100 mm. Two regions of interest (ROIs) were defined for each root (left and right are separate) above and below the DRG for tractography (example in **Figure 3B**) and named after that root. Tractography through each pair of ROIs was terminated once 10,000 tracts were identified to keep consistency in metric comparison. Sporadic fiber tracts and those that do not reach the spinal cord were pruned to obtain the final tracts. These tracts were also named after the root level it corresponds to. We then saved these tracts as .tt files and also extracted the coordinates (x, y, z) of these tracts into text files that were later used in the computational models.

#### Quantitative Diffusion and Tractography Metrics

In order to quantify diffusion and tractography results we computed several metrics including fractional anisotropy (FA), mean diffusivity (MD), tract length, tract diameter, and tract volume. FA is the fraction of diffusion that is direction-dependent where values vary from zero to one with zero corresponding to unrestricted diffusion in all directions, while one corresponds to diffusion along a single axis. MD describes the rotationally invariant magnitude of water diffusion within tissue and relates to the total amount of diffusion in a voxel. Tract length, diameter, and volume refer to the length, diameter, and volume of the diffusion based fiber tracts ^96^. These metrics were all exported from DSI Studio after tractography had been completed.

#### Optimization of diffusion directions

In DSI studio, the b-table for the 61 diffusion direction acquisition of C01-LS was imported. This b-table retains the same order as it was generated from the Emmanuel Caruyer online q-space sampling protocol application ^53^, which keeps uniform sampling distribution with each direction it adds. Thus, to simulate tractography results from fewer diffusion directions, we sequentially removed one b-value at a time from the b-table. After removal of each b-value, a tensor reconstruction was conducted using the new b-table, and this was repeated until we only have 6 directions left in the b-table. The same ROIs were used, and quantitative metrics exported for each of the 55 tested subsets of DTI for tractography. To optimize the number of diffusion directions, we plotted the quantitative metric values for each root level against the number of diffusion directions used for fiber tractography. We identified 19 diffusion directions to be the minimum and optimal number for which FA, MD, tract length, tract diameter, and tract volume flattened and remained stable, indicating a plateau in fiber tract reconstruction quality.

### Morphometric Quantifications

#### MRI measurements

##### Segmentation based

we performed several measurements based on the full segmentation of the spinal cord using the CNN-based automatic segmentation algorithm. These segmentations were validated to be as precise as a human expert and are thus reliable and suitable for automated morphological quantifications. These quantifications were performed in MATLAB (The MathWorks Inc., Natick, Massachusetts) and included gray and white matter areas, epidural fat area and thickness, CSF area and thickness, as well as spinal cord width and height across the full length of the imaging field of view. Spinal cord width and height were measured both directly on selective T2-weighted slices and automatically calculated using the white matter segmentation masks, which gave compatible results at voxel size precision, further validating our segmentation-based quantification method. For area calculations, we calculated the sum of pixels in the segmentation mask for the respective tissue in each axial image slice and multiplied it by the in-plane pixel dimension (50 x 50 µm^2^). For thickness calculations, since the spinal cord is not perfectly straight thus not always centered in the image, we manually set the boundary lines for dorsal and lateral on each slice using the function *ginput()*. Then, we calculated the coordinate difference between pixels closest and furthest to the boundary line and converted it to distance by multiplying pixel width (50 µm).

##### Image based

other measurements, including dorsal/ventral root and rootlet diameter, number of rootlets, vertebral height, DREZ length, and root angles for each root level, vary a lot in orientation, shape, and location in the spine, and were therefore measured directly from the images by a spine image specialist in either FSLeyes or Fiji (ImageJ) ^97^. These measures were taken mostly from the axial slice and verified in sagittal and coronal views. The root angles were measured in the sagittal slices, and the vertebral height from the mid coronal slice. A schematic of each of these morphological measurements are illustrated in **Figure S6**.

##### DTI tractography measurements

With the DTI tractography, we can clearly visualize the DREZs as well as the spinal roots, and thus we could perform measurements within DSI studio. For DREZ length, by overlaying the axial T2w slice with the tractography, we counted the number of slices from top of each DREZ level to the bottom of that DREZ, then calculated the length based on the slice thickness. For root diameter, we cut the fiber tracts for each root right after the DRG and before the root splits into nerve branches using the “Cut by slice” tool in DSI studio (http://dsi-studio.labsolver.org, CUDA SM75 version Apr 7, 2022), we then exported the tract diameter metric as described in the *Diffusion Tensor Image Analysis* section above.

##### Dissection measurements

We performed measurements directly on two lumbosacral spinal tissues (C01-LS, C03-LS) through dissection as ground truth for imaging measurements. To do this, we first measured the vertebral heights between the vertebral discs with a caliper (0.01 mm resolution). Then, we performed full laminectomy until the vertebral foramina, removed any epidural fat, and carefully made a medial incision on the dura to expose the spinal cord and rootlets, where we measured the vertical distance between the rostral end of the DREZ of each root (beginning of the spinal segment) to where the root exited the vertebra (end of the vertebral segment). Following that, we detached the nerve roots from the muscles and vertebral foramen and removed the spinal cord and roots from the spinal column. On the isolated spinal cord, we measured root diameters just past the DRG, dorsal/ventral rootlet diameters, DREZ lengths, cord width and height with the caliper, and counted the number of rootlets at each spinal level.

### Dorsal Root Ganglion Cell Body Distribution

#### MRI Processing

Using the FSL software, the T2w MR images were segmented by locating the DRG, identified by an enlargement on the dorsal side of the spinal root with inhomogeneous intensity. In T2w MRI, brighter pixels indicate cell bodies while darker pixels correspond to axons of sensory afferents. The boundary of the DRG was thus defined as the most spinal and peripheral axial slices containing cell bodies, based on intensity observations. The segmented DRGs were exported as NIFTI files and imported into MATLAB (version R2023b) using the *niftiread()* function provided by the Image Processing Toolbox.

#### Histology Processing

After all imaging and measurements were completed on the samples, we dissected the DRGs (preserving a short segment of spinal root) from the spinal cord to perform histological staining as a ground truth for the DRG cell distribution analysis. This was done for the L6 DRGs of sample C03-LS. Upon dissection, we used a tissue marking dye (Cancer Diagnostics, Inc, Durham, NC) to highlight the anatomical lateral side of the DRG (blue), since information about laterality is often lost during sectioning. The DRG tissue was kept in 4% PFA for one week, then dehydrated and paraffin-embedded for subsequent axial sectioning (10 µm, transverse to the spinal root) and chemical staining. The embedding and sectioning were performed by the Neuropathology Lab in the Department of Pathology at the University of Pittsburgh. For the most mid-axial sections of the DRG (widest), we used Nissl/Luxol Fast Blue to stain the cell bodies and myelinated axons respectively, using standard techniques. Sections with substantial damage were excluded from the analysis.

The stained slides were imaged through a Nikon Optiphot-2 microscope equipped with Spot Insight 2MP color camera, in the brightfield setting, and with PlanApo4-0.2 lens for 4x magnification using Spot software version 4.6. We imported the images into ImageJ ^98^ (https://imagej.nih.gov/ij/, RRID: SCR 003070), rotated the images so the DRG is on the top and the ventral root on the bottom. Then we manually demarcated the neuronal cell bodies (in blue) and the DRG border (in red) with the ObjectJ plugin ^99^ (https://sils.fnwi.uva.nl/bcb/objectj/index.html), as in Ostrowski et al. 2017 ^60^, and exported the labeled images as PNGs. Finally, we imported these images into MATLAB in RGB colors and extracted pixels corresponding to cell bodies and DRG based on the color of the demarcation.

#### Cell Distribution Quantifications

For both MRI segmented and histology segmented DRGs, a polar transformation was applied to each slice to standardize the area and enable comparison between DRGs of different sizes and shapes. The transformation process is based on the method presented in Ostrowski et al. 2017 ^60^. Briefly, a centroid is defined as the origin, and each voxel is assigned a polar angle from 0° to 360° (0° corresponds to the positive horizontal axis). The distance of each voxel, within the DRG border, from the centroid is calculated and normalized by the distance of the DRG border from the centroid at the same angle. The coordinates of these voxels are then mapped onto a unit circle. Then, the circle is divided into 8 x 6 areas, delineated by 8 angular sectors and 6 rings. The selection of 8x6 areas is chosen to suit analysis comparing various regions. For MRI images, density is calculated based on the mean intensity of all the voxels within that area, while for histology sections, density is calculated by the percentage of voxels that are cell bodies within the area. A right mid-DRG slice of cat 3 at level L6 was selected for both techniques and compared. Three distinct analyses were then conducted: 1) comparison between the ventral and dorsal sides, 2) comparison between the inner, middle, and outer rings, 3) comparison between medial and lateral sides of the DRG.

### Computational Modeling

#### Complex spine model

We designed a computational framework to develop an anatomically and neurofunctionally realistic computational model of the cat lumbosacral spine. To achieve this, we exploited tissue contours (gray matter, white matter, dorsal roots and rootlets, ventral roots and rootlets, cerebral spinal fluid, and epidural fat) automatically segmented by the modified HarDNet. Prior to the extraction of 3D volumes, we imported micro-CT vertebrae segmentations from thresholding and performed manual correction of the identified tissue boundaries using iSeg (ZMT), including re-labeling of misclassified pixels and smoothing of contours. We also added a 50 µm thick shell around the CSF to represent the dura mater. Post-processed segmentations were imported in Sim4Life 7.0 (IT’IS Foundation/ZMT Zurich MedTech AG), a multi-physics simulation platform. We extracted 3D volumes using the *Surface Extraction* tool, and added a cylinder of radius 20 mm around the spine to mimic muscular and connective tissue and to fill any gaps in the spinal volumes; we generated an additional saline cylinder of radius 25 mm. Using a structured mesh engine, we generated cubic voxels with varying resolution across different compartments, with roots having the highest resolution. Tissues were assigned isotropic conductivity values (0.23 S/m for gray matter, 1.7 S/m for CSF, 0.04 S/m for epidural fat, 0.02 S/m for bone, 2 S/m for saline, 0.2 S/m for muscle/connective tissue) from literature ^65,100,101^. For the dorsal and ventral roots and rootlets, as well as the white matter, we calculated anisotropic conductivity maps by aligning conductivity tensors (conductivities were set to 0.083 S/m transverse and 0.6 S/m longitudinal to the tensor) to the tissue diffusion gradient as described in Rowald et al., 2022 ^64^. Finally, we functionalized the model with realistic axon trajectories derived from the dMRI and populated each spinal level (L6 to S3), location (left and right) and side (dorsal and ventral) with 25 neurons, resulting in a total of 500 neurons in each model.

#### Canonical spine model

We designed a canonical model, adopting commonly reported methods without the use of a full set of high-resolution imaging. The dimensions of gray and white matter were manually segmented from 11 MRI slices evenly distributed across the rostro-caudal extent of the lumbosacral spine, and interpolated to obtain 3D volumes in Solidworks (Version 2022). We derived the dimensions of the CSF and epidural fat from a previously published human lumbosacral spinal cord model ^102^, scaled to match the cat dimensions.

The vertebrae structure was simplified as a cylindrical shell around the spine. No dura, dorsal or ventral roots and rootlets volumes were included in the model. To design simplified axonal trajectories, we manually inserted five bundles of five rootlets for each segment (L6-S3), location and side. Geometrical dimensions about spinal segment length of the cat lumbosacral spinal cord were based on data from previous literature ^79^, while the roots angle (with respect to the spinal cord midline) were derived from anatomical measurements on the average of our cat spine samples (C01-LS, C03-LS, C06-LS).

#### Simulation of neural recruitment

For both complex and canonical spine models, we modeled stimulating electrodes as 0.291 mm x 1.0 mm x0.05 mm silicone volumes encapsulating active contact, placed under the L6 vertebra.

An electromagnetic field was generated by applying a biphasic pulse of 1 V with a pulse width of 0.2 ms in each phase on the active site at each vertebral level. We imposed a Dirichlet boundary condition (0 V) on the outer surface of the saline volume. The resulting field was then applied to the neural paths generated manually or via DTI, which were functionalized as NEURON models using the NEURON environment integrated in Sim4Life 7.0.

We assigned uniform axon diameters of 12 µm to ensure that analyses captured differences solely due to the geometric differences between trajectories. Neurons in the dorsal and ventral roots were implemented in Sim4Life as the Motor and Sensory MRG model ^103,104^.The electromagnetic field was then scaled linearly to determine recruitment threshold of the NEURON axons using the built-in Sim4Life Titration tool. We compared the population recruitment amplitude of afferent and efferent axons for both the canonical and realistic model. To estimate simulated selectivity, we normalized the recruitment amplitude for each model by dividing each axon’s recruitment threshold for the maximum recruitment threshold across afferents and efferents. Finally, we computed the normalized threshold difference by comparing the normalized afferent-efferent threshold at each root level (L6 to S3) for the canonical and realistic model.

## STATISTICAL ANALYSIS

Statistical comparisons were performed for several analyses in this manuscript. Since sample sizes are small, we cannot assume normal distribution, and groups for comparison can all be considered independent, we therefore used non-parametric tests chosen based on the number of groups being compared. All statistical comparisons were performed in MATLAB (The MathWorks Inc., Natick, Massachusetts) versions 2019b-2023b. For DRG cell distribution analysis, we used the Mann-Whitney U-test (a.k.a Wilcoxon Rank Sum test) to compare between the dorsal and ventral regions, as well as between the medial and lateral sides, while for the ring analysis (outer, middle, inner), we used the Kruskal-Wallis test with a Bonferroni correction. Similarly, for the neural network input channel analysis, to compare between five different input channel combinations, we used the Kruskal-Wallis test with Bonferroni correction. Finally, for the computational neuron recruitment analysis, we used the Kruskal-Wallis test with Bonferroni correction for absolute recruitment amplitude comparisons between afferents and efferents of and across the canonical and realistic models. For comparison between the normalized threshold differences of canonical and realistic models, we used the Mann-Whitney U-test (a.k.a Wilcoxon Rank Sum test). Significant differences between groups are indicated with stars in the corresponding plots: * for p-value < 0.05, ** for p-value < 0.01, and *** for p-value < 0.001. Additional details on the number of data points (n) used for analysis and dispersion measures displayed in plots can be found within specific figure captions.

## DATA AND CODE AVAILABILITY

All imaging data acquired for this study and the protocols will be available on The Common Fund’s Stimulating Peripheral Activity to Relieve Conditions (SPARC) program data sharing platform at sparc.science and protocols.io upon publication.

## AUTHOR CONTRIBUTIONS

EP and TKH conceived and supervised the study. RAG, EP, TKH, LEF, MC, EN secured funding and resources. LL, EP, TKH designed the imaging protocols and collected the data. DJS performed micro-CT imaging. LL, UA, CR, AD, MDB performed dissection and measurements of the cat spine samples. LL processed and analyzed the images, and LL, UA, CR performed the manual segmentations. AF designed and LL assisted with optimization of the automatic segmentation algorithm. AD built and MJ and TN assisted with the canonical and complex spine FEM & Neuron models for simulation comparisons. MDB and LEF secured and processed the cadaver spine sample. RM, LL, and CLK performed DRG histology and cell distribution analysis. The manuscript was written by LL, EP, and TKH with assistance from AD and RM. All authors contributed to its revision.

## ACKNOWLEDGMENTS

We thank Tyler Simpson for designing and building containers for the human cadaver sample, Dylan Beam for assisting in the cadaver sample dissections, Rachel Pitzer for managing all the materials involved in this project, and Max Novelli, and Sam Ward for data curation and IT support, all from the Rehab Neural Engineering Labs. We also thank Lesley Foley for assistance with MRI studies, and Jinho Kim and Kelly Puglisi for their help with histological processing and imaging. Additionally, we thank Dr. Frank Yeh and Fang Liu for their helpful discussions and valuable support with image processing. Magnetic Resonance Imaging was performed in the Advanced Imaging Center (RRID:SCR_025139) at the University of Pittsburgh, supported in part by the office of Senior Vice Chancellor for Health Sciences. Finally, we would also like to thank the deceased and their family for generously donating the human spine tissue used in this work. This work was supported by the Common Fund’s Stimulating Peripheral Activity to Relieve Conditions (SPARC) of the National Institutes of Health with award number OT2OD030537.

## SUPPLEMENTAL INFORMATION

### Supplementary Tables

**Supplementary Table 1.**
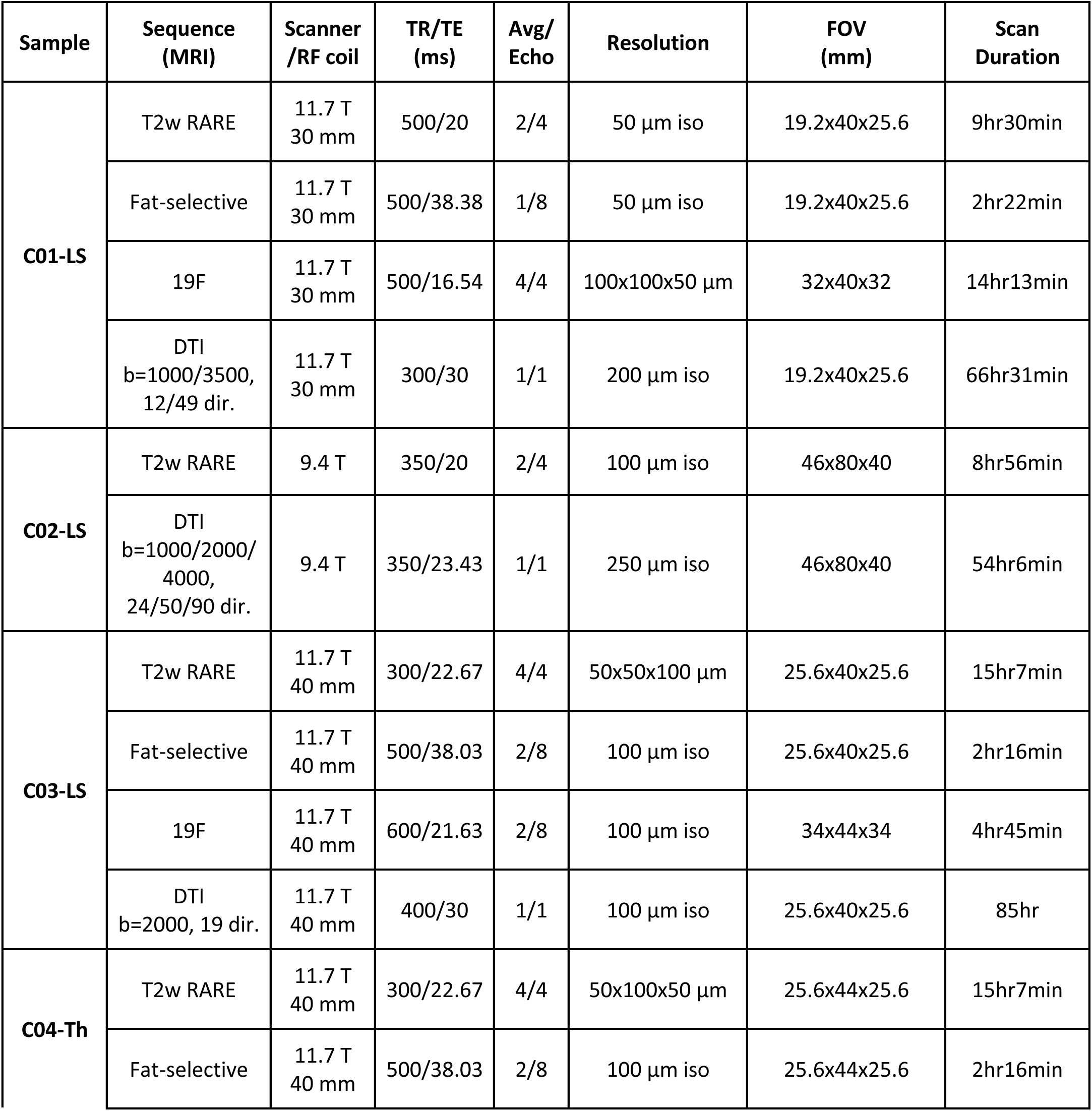
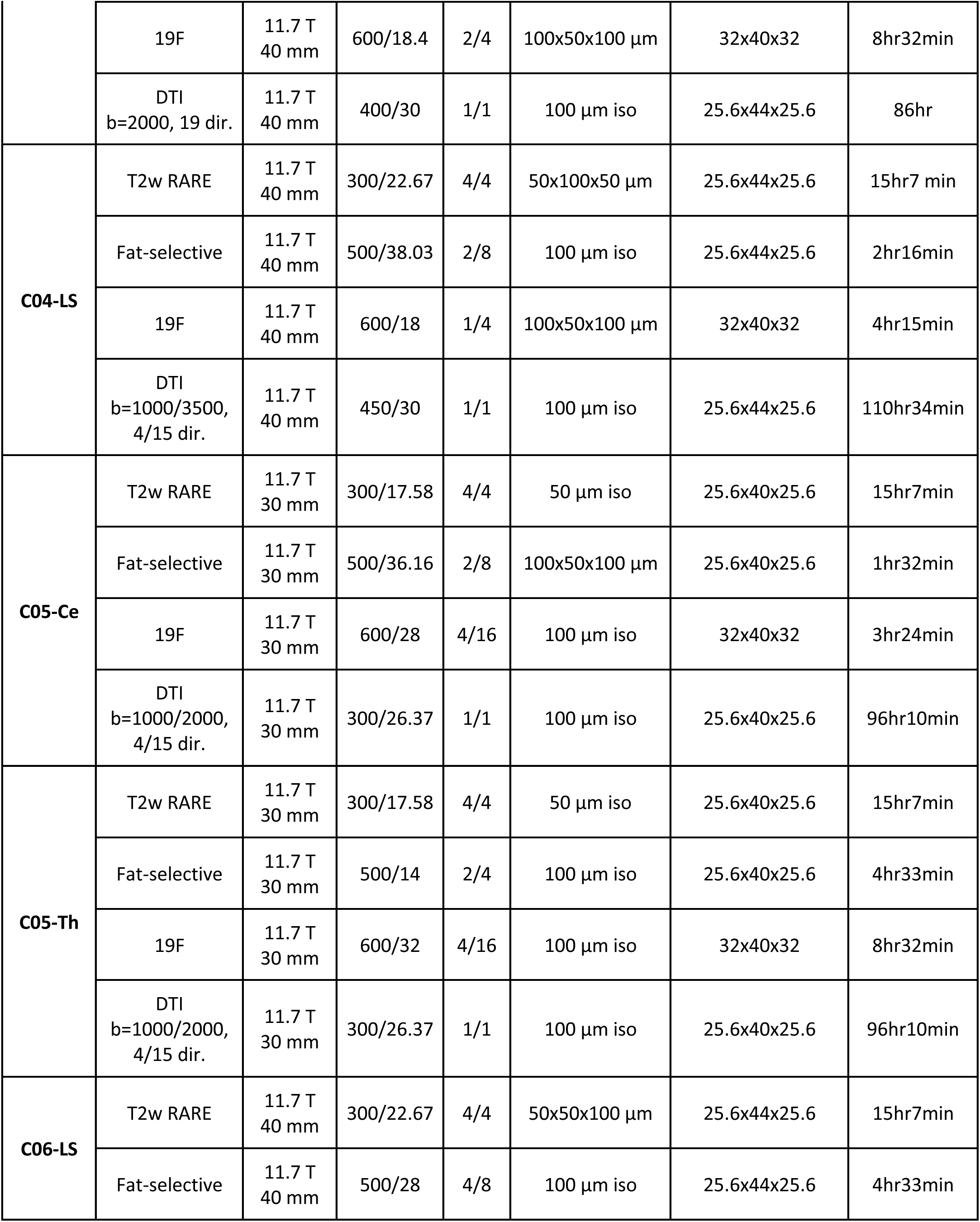
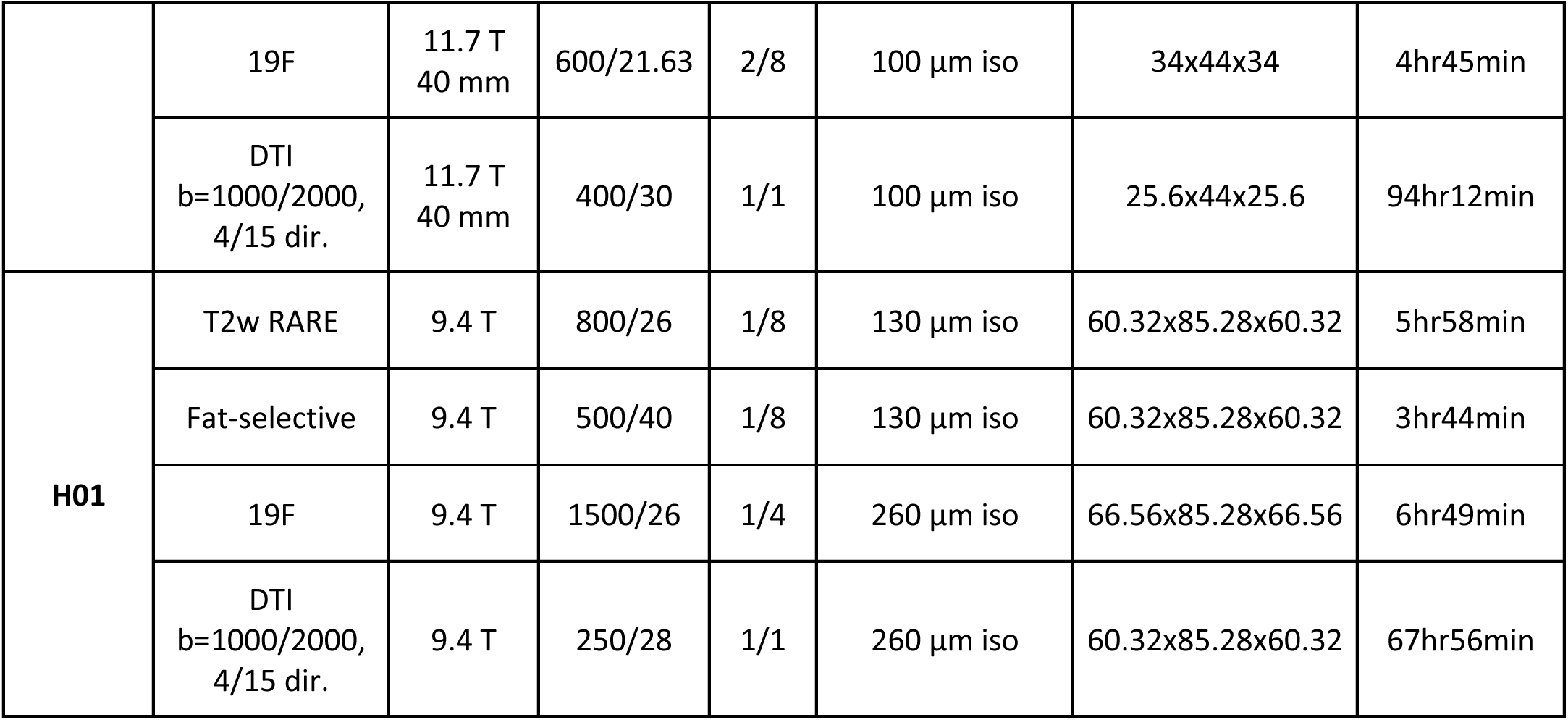
MRI scan sequences and parameters for all samples. All sequences were 3D. Fat-selective MRI was collected with narrow excitation at −3.5 ppm, 19F MRI was collected at 470 MHz with the carrier on the Fomblin major resonance peak. FOV: field of view; TR/TE: repetition time/echo time; RF: radio frequency.

| Sample | Sequence (MRI) | Scanner /RF coil | TR/TE (ms) | Avg/ Echo | Resolution | FOV (mm) | Scan Duration |
| --- | --- | --- | --- | --- | --- | --- | --- |
| **C01-LS** | T2w RARE | 11.7 T<br>30 mm | 500/20 | 2/4 | 50 μm iso | 19.2x40x25.6 | 9hr30min |
| | Fat-selective | 11.7 T<br>30 mm | 500/38.38 | 1/8 | 50 μm iso | 19.2x40x25.6 | 2hr22min |
| | 19F | 11.7 T<br>30 mm | 500/16.54 | 4/4 | 100x100x50 μm | 32x40x32 | 14hr13min |
| | DTI<br>b=1000/3500,<br>12/49 dir. | 11.7 T<br>30 mm | 300/30 | 1/1 | 200 μm iso | 19.2x40x25.6 | 66hr31min |
| **C02-LS** | T2w RARE | 9.4 T | 350/20 | 2/4 | 100 μm iso | 46x80x40 | 8hr56min |
| | DTI<br>b=1000/2000/<br>4000,<br>24/50/90 dir. | 9.4 T | 350/23.43 | 1/1 | 250 μm iso | 46x80x40 | 54hr6min |
| **C03-LS** | T2w RARE | 11.7 T<br>40 mm | 300/22.67 | 4/4 | 50x50x100 μm | 25.6x40x25.6 | 15hr7min |
| | Fat-selective | 11.7 T<br>40 mm | 500/38.03 | 2/8 | 100 μm iso | 25.6x40x25.6 | 2hr16min |
| | 19F | 11.7 T<br>40 mm | 600/21.63 | 2/8 | 100 μm iso | 34x44x34 | 4hr45min |
| | DTI<br>b=2000, 19 dir. | 11.7 T<br>40 mm | 400/30 | 1/1 | 100 μm iso | 25.6x40x25.6 | 85hr |
| **C04-Th** | T2w RARE | 11.7 T<br>40 mm | 300/22.67 | 4/4 | 50x100x50 μm | 25.6x44x25.6 | 15hr7min |
| | Fat-selective | 11.7 T<br>40 mm | 500/38.03 | 2/8 | 100 μm iso | 25.6x44x25.6 | 2hr16min |
| | 19F | 11.7 T<br>40 mm | 600/18.4 | 2/4 | 100x50x100 μm | 32x40x32 | 8hr32min |
| | DTI<br>b=2000, 19 dir. | 11.7 T<br>40 mm | 400/30 | 1/1 | 100 μm iso | 25.6x44x25.6 | 86hr |
| **C04-LS** | T2w RARE | 11.7 T<br>40 mm | 300/22.67 | 4/4 | 50x100x50 μm | 25.6x44x25.6 | 15hr7 min |
| | Fat-selective | 11.7 T<br>40 mm | 500/38.03 | 2/8 | 100 μm iso | 25.6x44x25.6 | 2hr16min |
| | 19F | 11.7 T<br>40 mm | 600/18 | 1/4 | 100x50x100 μm | 32x40x32 | 4hr15min |
| | DTI<br>b=1000/3500,<br>4/15 dir. | 11.7 T<br>40 mm | 450/30 | 1/1 | 100 μm iso | 25.6x44x25.6 | 110hr34min |
| **C05-Ce** | T2w RARE | 11.7 T<br>30 mm | 300/17.58 | 4/4 | 50 μm iso | 25.6x40x25.6 | 15hr7min |
| | Fat-selective | 11.7 T<br>30 mm | 500/36.16 | 2/8 | 100x50x100 μm | 25.6x40x25.6 | 1hr32min |
| | 19F | 11.7 T<br>30 mm | 600/28 | 4/16 | 100 μm iso | 32x40x32 | 3hr24min |
| | DTI<br>b=1000/2000,<br>4/15 dir. | 11.7 T<br>30 mm | 300/26.37 | 1/1 | 100 μm iso | 25.6x40x25.6 | 96hr10min |
| **C05-Th** | T2w RARE | 11.7 T<br>30 mm | 300/17.58 | 4/4 | 50 μm iso | 25.6x40x25.6 | 15hr7min |
| | Fat-selective | 11.7 T<br>30 mm | 500/14 | 2/4 | 100 μm iso | 25.6x40x25.6 | 4hr33min |
| | 19F | 11.7 T<br>30 mm | 600/32 | 4/16 | 100 μm iso | 32x40x32 | 8hr32min |
| | DTI<br>b=1000/2000,<br>4/15 dir. | 11.7 T<br>30 mm | 300/26.37 | 1/1 | 100 μm iso | 25.6x40x25.6 | 96hr10min |
| **C06-LS** | T2w RARE | 11.7 T<br>40 mm | 300/22.67 | 4/4 | 50x50x100 μm | 25.6x44x25.6 | 15hr7min |
| | Fat-selective | 11.7 T<br>40 mm | 500/28 | 4/8 | 100 μm iso | 25.6x44x25.6 | 4hr33min |
| | 19F | 11.7 T<br>40 mm | 600/21.63 | 2/8 | 100 μm iso | 34x44x34 | 4hr45min |
| | DTI<br>b=1000/2000,<br>4/15 dir. | 11.7 T<br>40 mm | 400/30 | 1/1 | 100 μm iso | 25.6x44x25.6 | 94hr12min |
| **H01** | T2w RARE | 9.4 T | 800/26 | 1/8 | 130 μm iso | 60.32x85.28x60.32 | 5hr58min |
| | Fat-selective | 9.4 T | 500/40 | 1/8 | 130 μm iso | 60.32x85.28x60.32 | 3hr44min |
| | 19F | 9.4 T | 1500/26 | 1/4 | 260 μm iso | 66.56x85.28x66.56 | 6hr49min |
| | DTI<br>b=1000/2000,<br>4/15 dir. | 9.4 T | 250/28 | 1/1 | 260 μm iso | 60.32x85.28x60.32 | 67hr56min |

**Supplementary Table 2.**
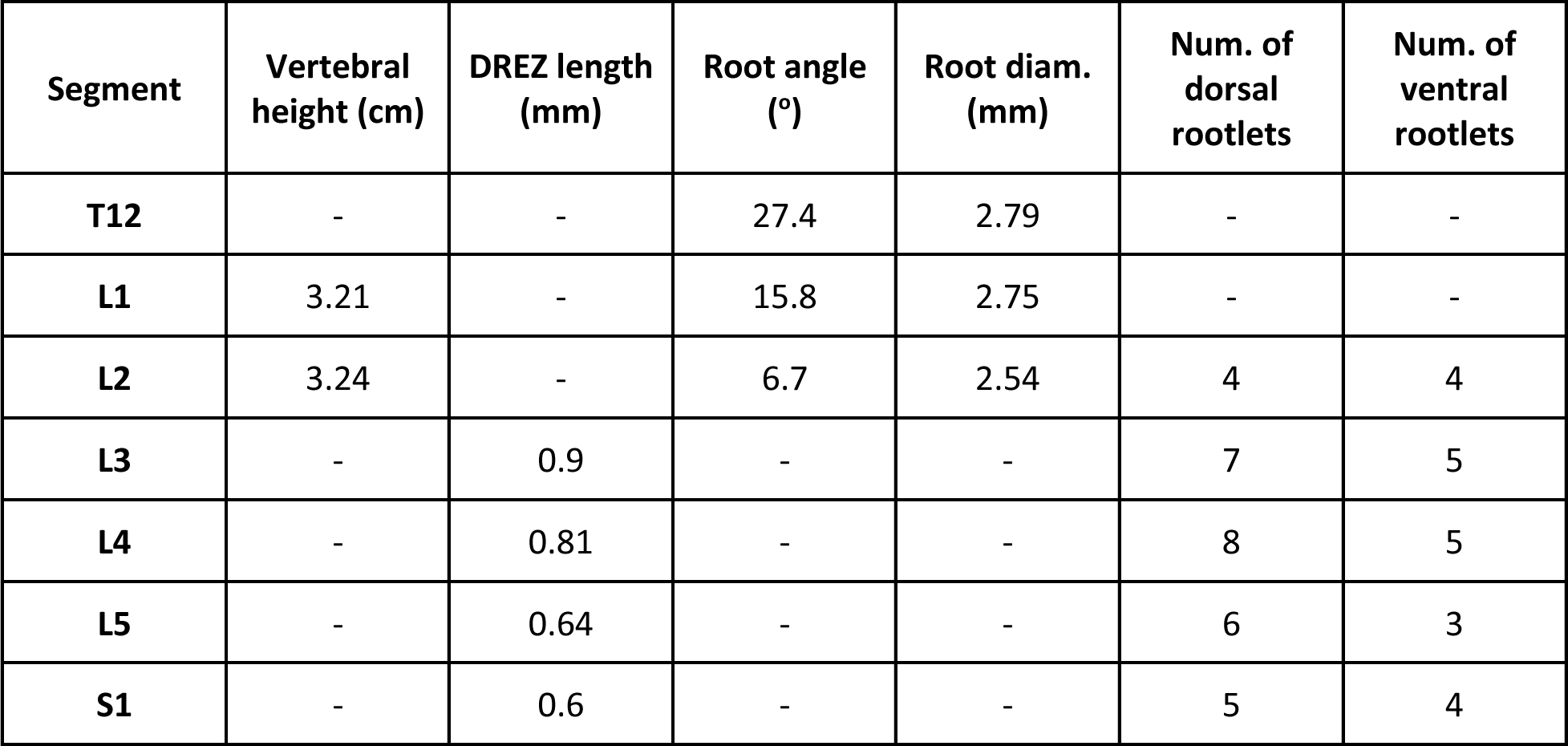
Gross spinal cord, root and rootlet morphometric measurements through MRI for human cadaver sample H01. Vertebral levels and spinal levels are significantly shifted in the human thoracic-lumbar spine, thus the segments measured for vertebral and spinal landmarks are also shifted.

**Supplementary Table 3.** Anatomical measurements obtainable by manual dissection versus SpIC3D. -: cannot be measured, o: can be measured. CSF: corticospinal fluid space; gm/wm: gray matter and white matter; DREZ: dorsal root entry zone.

|  |  | vertebra | epidural fat | CSF | vertebral venous plexus | spinal cord | gm/wm | root | rootlet | DREZ | root angle |
| --- | --- | --- | --- | --- | --- | --- | --- | --- | --- | --- | --- |
| Dissection |  | o | - | - | - | o | - | o | o | o | - |
| SpIC3D | T2/19F | o | o | o | o | o | o | o | o | o | o |
|  | DTI tract. | - | - | - | - | o | - | o | o | o | o |
|  | CT | o | - | - | - | - | - | - | - | - | - |

### Supplementary Figures

**Figure S1.**
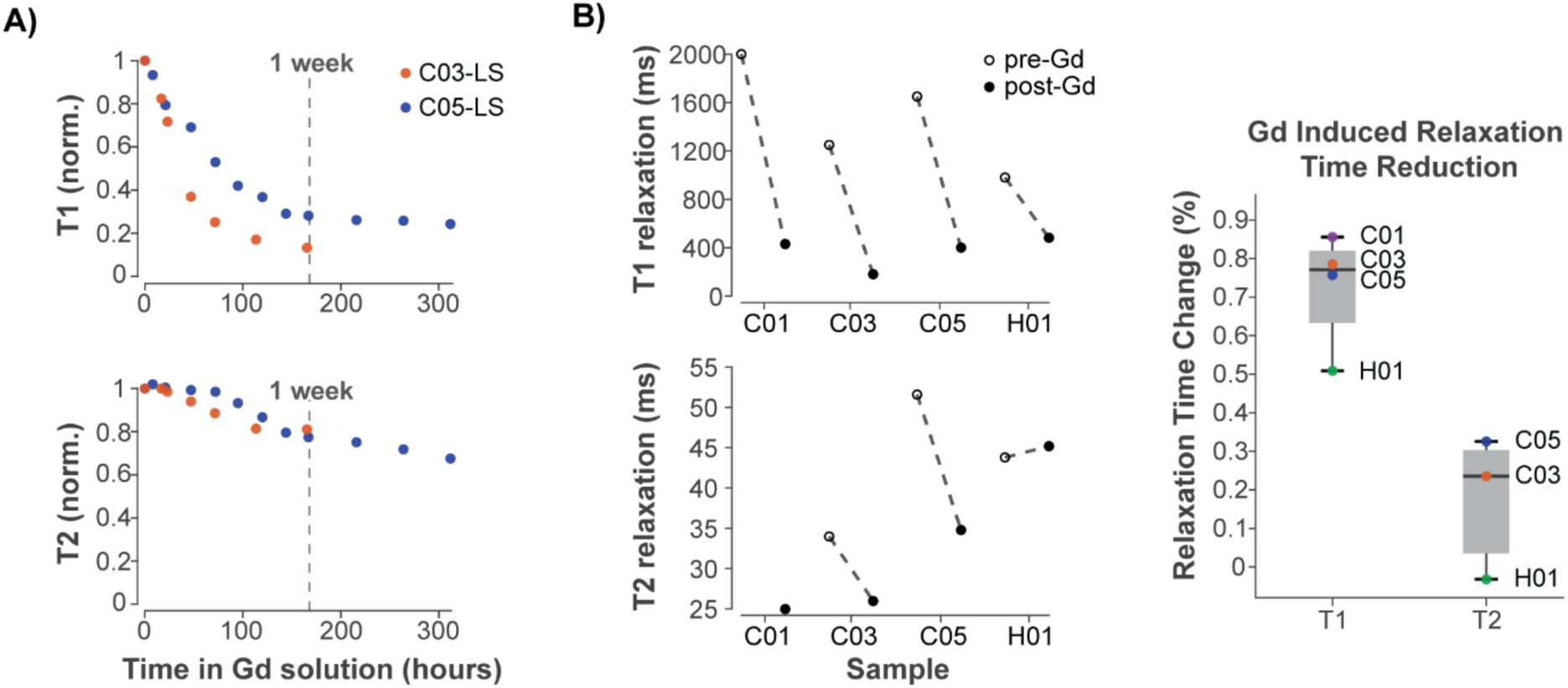
Relaxation time changes over time are consistent across samples, Related to Figure 1. **(A)** Normalized T1 (*top*) and T2 (*bottom*) relaxation change in 0.2% Gd overtime for the C03 and C05 lumbosacral cord, showing that the change pattern is consistent across animals for the same spinal level, i.e., lumbosacral. **(B)** *Left*: absolute value of T1 (*top*) and T2 (*bottom*) relaxation times pre and post (C01-LS: 2 weeks, C03-LS: 2 weeks, C05-LS: 1 week, H01: 2 weeks) submersion in Gadolinium for 4 samples, including human cord (H01). T1 values are consistent across samples and species. T2 changes are more variable but consistently small (<55 ms) for all samples. *Right*: box plot of percent relaxation time changes for all samples. For all boxplots, the whiskers extend to the maximum and minimum. Central line, top, and bottom of the box represent median,75th percentile, and 25th percentile, respectively.

**Figure S2.**
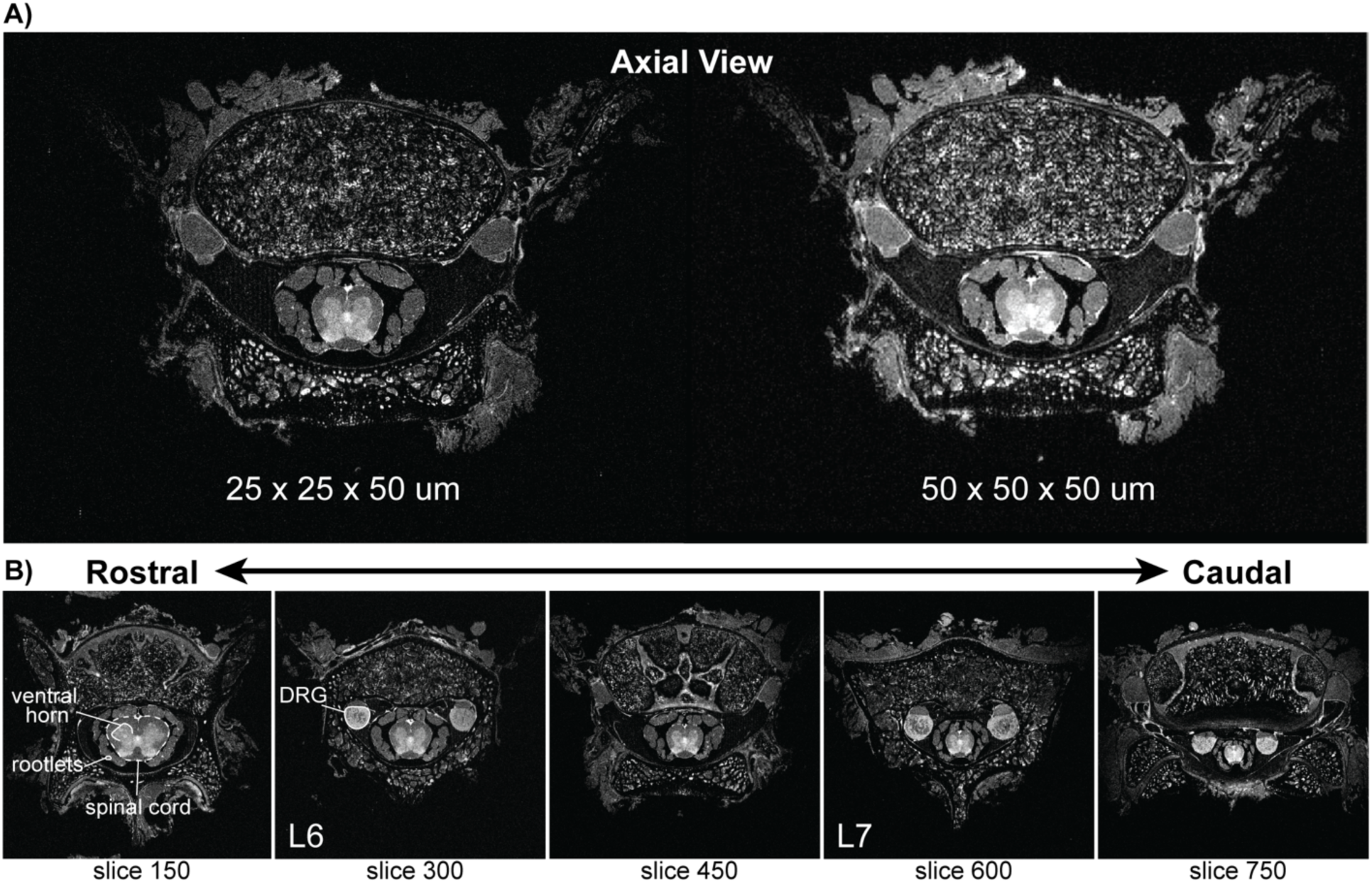
Examples of 25 µm T2 images, Related to Figure 2. **(A)** Comparison of the same axial slice in a 25 µm in-plane and a 50 µm in-plan resolution MRI of sample C01-LS. Both images show good tissue contrast. **(B)** Multiple axial MRI slices from the 25 µm acquisition showing rostro-caudal morphology changes, such as reduced ventral horn width, gradually decreasing spinal cord diameter and rootlet size, and different DRG sizes with L7 being the largest.

**Figure S3.**
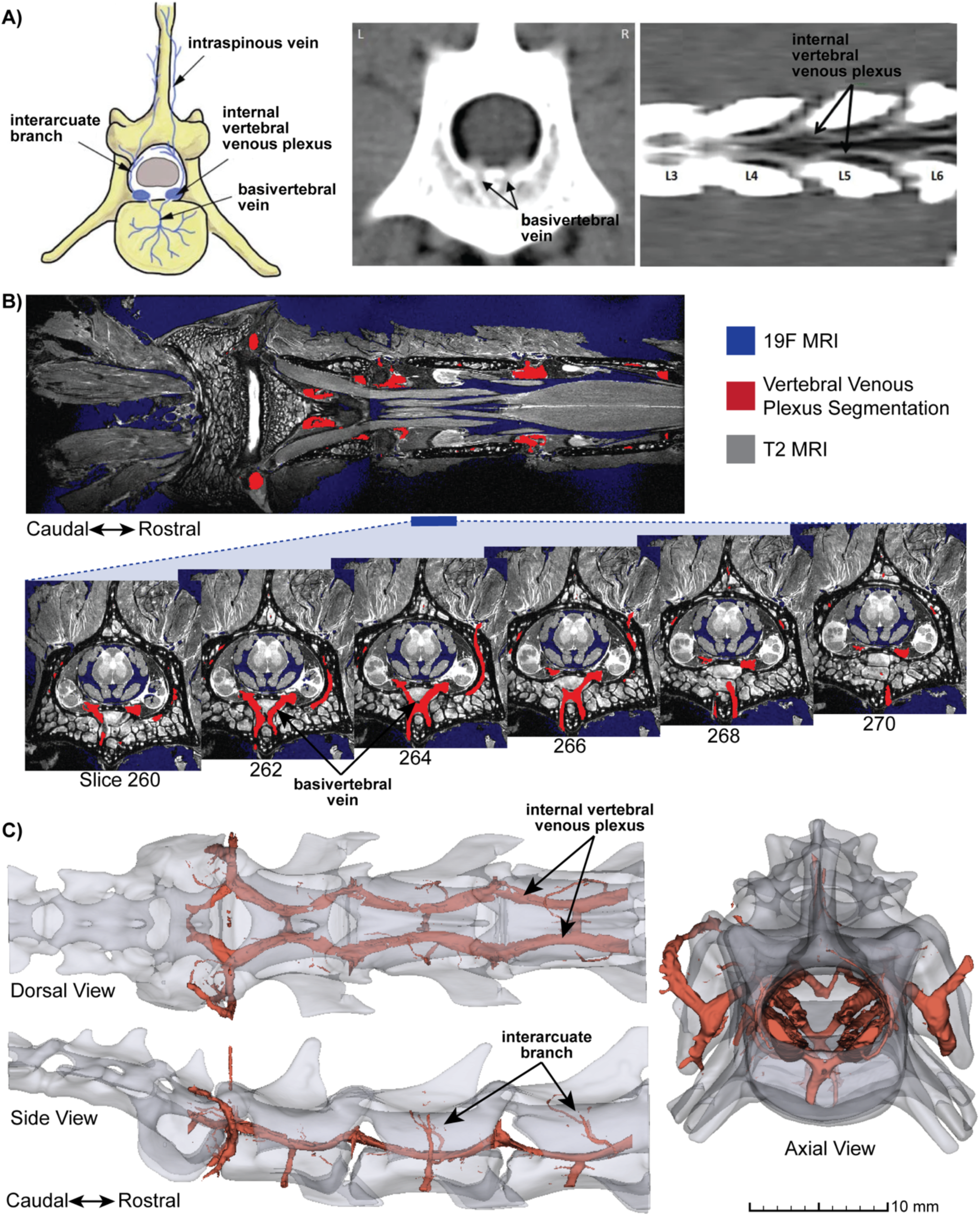
Accurate reconstruction of lumbar vertebral venous plexus, Related to Figure 2. **(A)** Schematic representation of some components of the vertebral venous plexus anatomy, extracted from an axial and a coronal view of a contrast CT venography of the canine lumbar spine. Adapted from Ariete et al., 2021 ^52^. **(B)** Coronal view of segmented vertebral venous plexus vessels captured by the 19F Fomblin MRI, overlaid on T2-weighted MRI. Selected axial slices from the middle of the L7 vertebrae area show the basivertebral vein coursing through the vertebral body. **(C)** 3D reconstruction of the segmented vertebral venous plexus in relation to the vertebrae, in dorsal, side, and top axial views. In addition to the ‘X’ shaped basivertebral vein, the thick internal vertebral venous plexus and interarcuate branches are also clearly visible.

**Figure S4.**
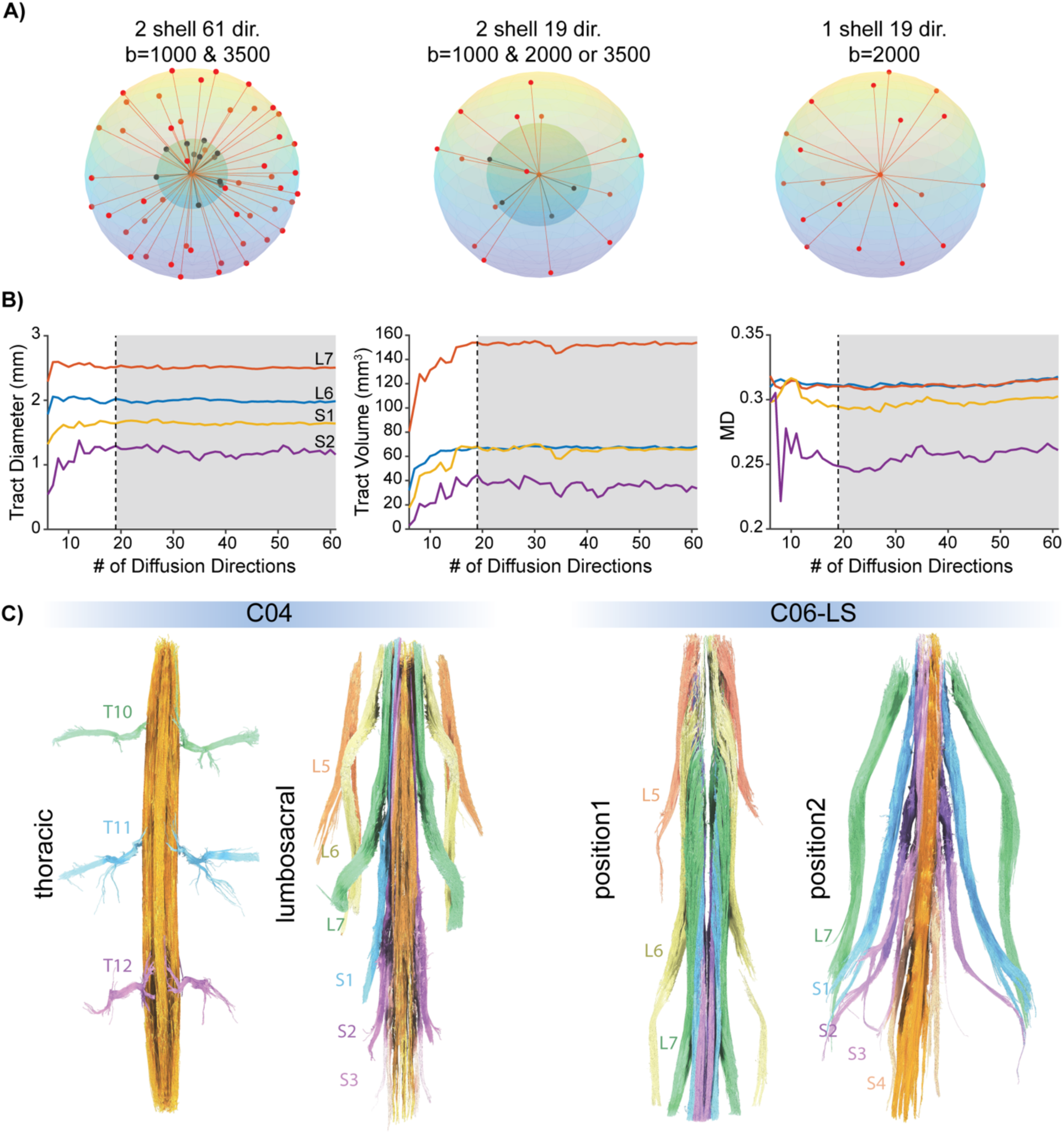
Additional single/dual shell 19dir DTI acquisition quality metrics and samples, Related to Figure 3. **(A)** Schematic representation of diffusion directions and corresponding shells used for DTI acquisitions. Diffusion direction schemes were generated with the Emmanuel Caruyer online q-space sampling protocol application ^53^ (http://www.emmanuelcaruyer.com/q-space-sampling.php) *Left*: 61 directions 2 shell setup used for sample C01- LS. *Middle*: 19 directions 2 shell setup used for samples C04-LS, C05-Ce, C05-Th, C06-LS. *Right*: 19 directions single shell setup used for samples C03-LS and C04-Th. **(B)** Assessment of tractography quality with varying number of diffusion directions (6-61 dir); convergence of tract diameter, tract volume and mean diffusivity (MD) measurements is observed for 15 directions and more. The dashed lines mark 19 directions, which was used to image all samples in this paper except C01-LS (i.e., the sample used for optimizing the DTI directions). **(C)** DTI tractography results for samples C04-Th, C04-LS, C06-LS position 1 (L5-L7) and position 2 (L7-S4+), with tracks colored according to root level. C04-LS is missing the right S1 root tract. The root tract was physically missing, as confirmed through dissection.

**Figure S5.**
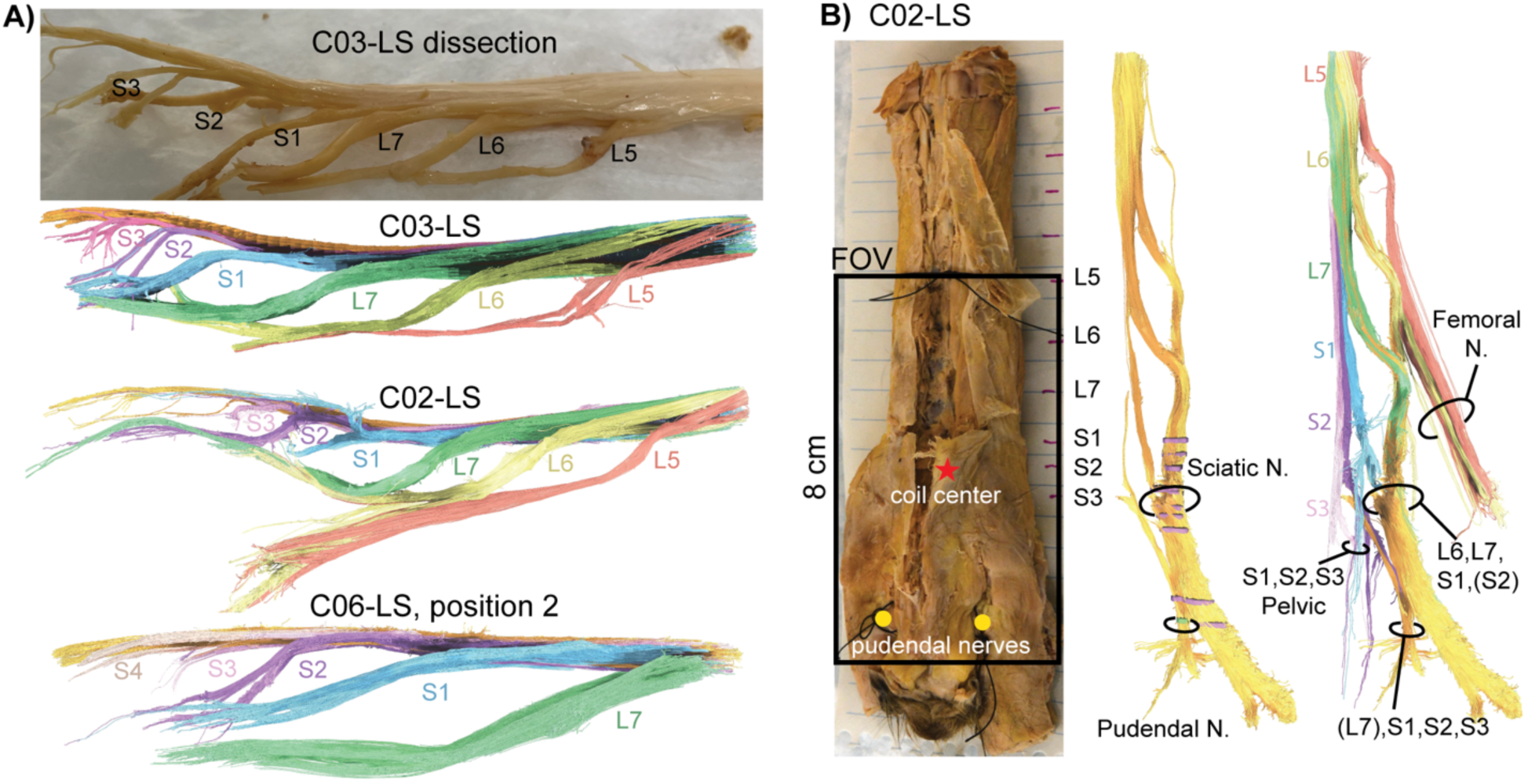
Identification of nerve branches and corresponding roots, Related to Figure 4. **(A)** Additional examples of DTI fiber tracking of the sacral plexus from multiple lumbosacral samples (C03-LS, C02- LS, C06-LS position 2). The C03-LS cord was carefully dissected to preserve its natural curvature and root orientations, and the in-situ tractography presents very high similarity with the dissected cord. Note: Lengths between samples are not to scale, but the color representing each root level is consistent. **(B)** *Left*: Extracted C02-LS sample preserving sciatic and pudendal nerves (i.e., larger sample). Red star: coil center and center of 8 cm FOV (marked by rectangle). Yellow dots: bilateral pudendal nerves marked with sutures. *Middle*: fiber tractography with ROIs drawn at the Sciatic N. and Pudendal N., showing contribution of fibers from each level of the spine. *Right*: fiber tractography with ROIs drawn around each root level, showing formation of the Femoral N., Sciatic N., Pudendal N., and possibly Pelvic N.

**Figure S6.**
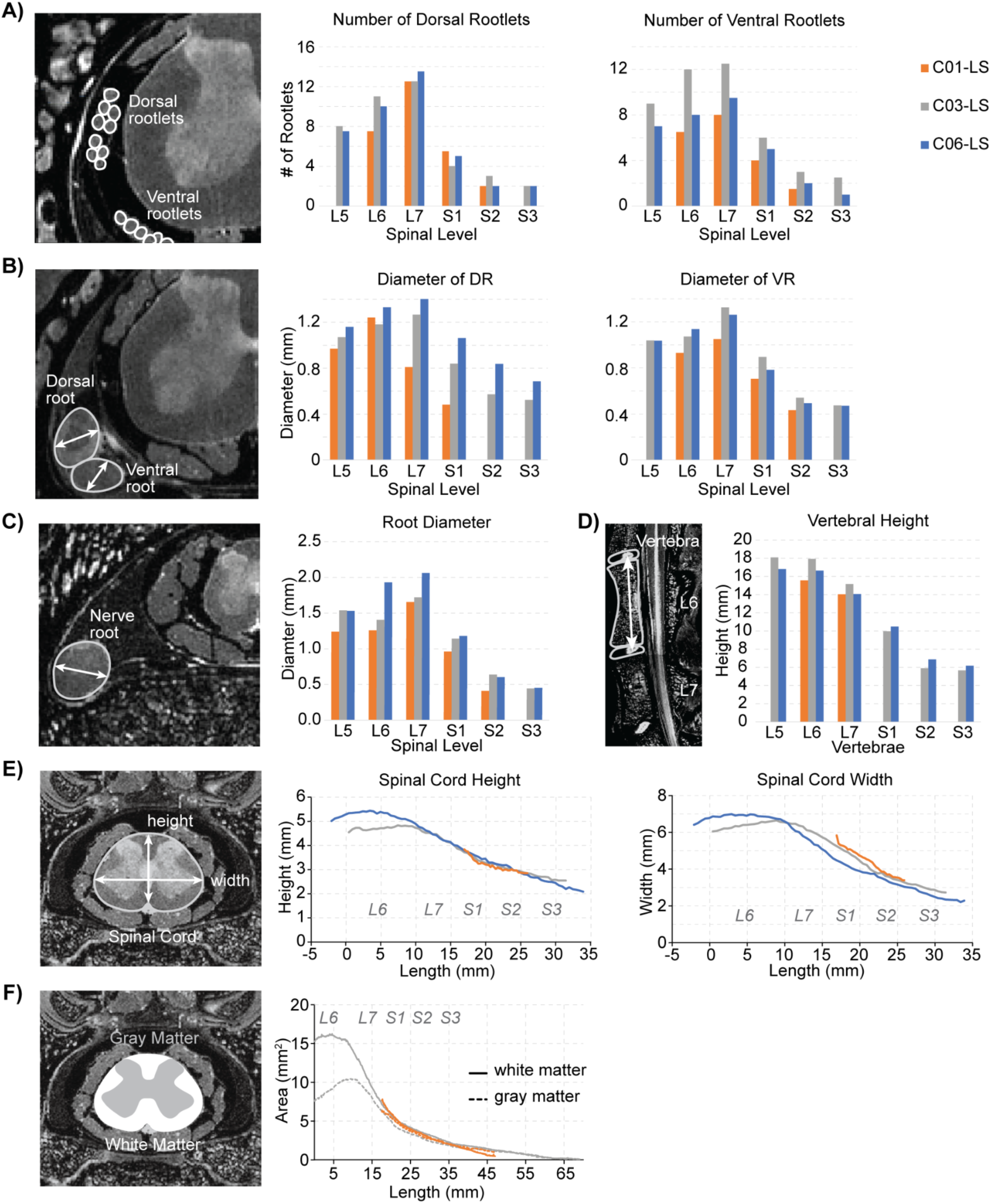
MRI-based spinal morphometry of cat lumbosacral samples, Related to Figures 2 and 4. **(A)** Dorsal and ventral rootlets visualized in axial MRI slices, as well as bar plots showing the number of dorsal and ventral rootlets for each spinal level for cat lumbosacral samples C01-LS, C03-LS, and C06-LS. Since C01-LS is shorter than the other samples, measurements from some levels are missing. **(B)** Dorsal root and ventral root in the axial MRI, as well as bar plots comparing their diameters across the samples from (A). **(C)** Nerve root past the DRG in the axial MRI, with bar plots of its diameter-dependence on spinal level for the different samples. **(D)** Vertebra in a sagittal MRI slice, and bar plots of the vertebra heights for the different samples. **(E), (F)** Spinal cord height and width, as well as gray and white matter areas along the length of the imaged samples, respectively. The L6, L7, S1, S2, S3 labels correspond to the respective spinal levels. C06-LS did not have good enough gray vs. white matter contrast to quantify the areas.

**Figure S7.**
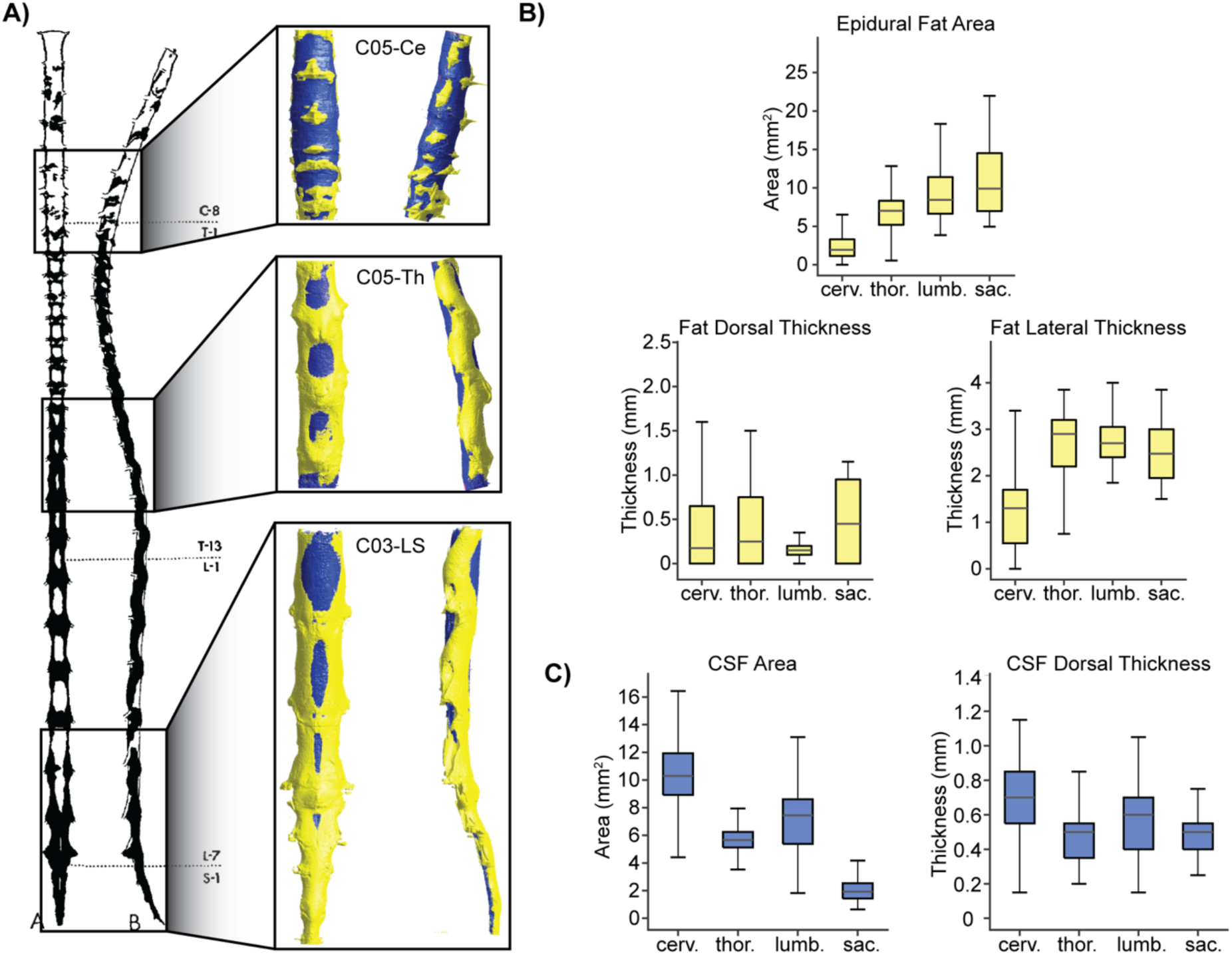
Epidural fat and CSF distribution comparisons across spine levels, Related to Figure 8. **(A)** Illustration of epidural fat distribution in the cat spinal cord from Ramsey et al. 1959 ^73^. Insets show epidural fat (yellow) reconstructed from SpIC3D of samples C05-Ce, C05-Th, and C03-LS, which closely resemble the illustration. **(B)** Boxplots of epidural fat area, dorsal thickness, and lateral thickness compared between the cervical (C05-Ce, n=800 slices), thoracic (C05-Th, n=800 slices), lumbar (C01-LS, n=610 slices; C03-LS n=490 slices), and sacral segments (C01-LS, n=114 slices; C03-LS, n=200 slices). **(C)** Boxplots of cerebro-spinal fluid (CSF) area and dorsal thickness across cervical, thoracic, lumbar, and sacral segments. For all boxplots, the whiskers extend to the maximum and minimum, excluding outliers. We did not report lateral thickness for CSF because the trends were comparable to dorsal thickness. Central line, top, and bottom of the box represent median, 75th percentile, and 25th percentile, respectively.

